# Feedback control of morphogen gradient scale

**DOI:** 10.1101/824292

**Authors:** Yilun Zhu, Yuchi Qiu, Weitao Chen, Qing Nie, Arthur D. Lander

## Abstract

Gradients of the morphogen decapentaplegic (Dpp) pattern *Drosophila* wing imaginal discs, establishing gene expression boundaries at specific locations. As discs grow, Dpp gradients expand, keeping relative boundary positions approximately stationary. Such scaling fails in mutants for *Pentagone* (*pent*), a gene repressed by Dpp that encodes a diffusible protein that expands Dpp gradients. Although these properties fit a recent mathematical model of automatic gradient scaling, we show here that Pent lacks a property essential to that model—the ability to spread with minimal loss throughout the morphogen field. Instead, Pent’s actions appear confined to within a few cell diameters of its site of synthesis, and can be phenocopied by manipulating non-diffusible targets of Pent strictly within the Pent expression domain. Through genetic manipulation and mathematical modeling we develop an alternative model of scaling, driven by feedback down-regulation of Dpp receptors and co-receptors. Among the model’s predictions is a size limit beyond which scaling fails—something we observe directly in wing discs.

## INTRODUCTION

During development, gradients of secreted morphogens convey positional information that enables cells to behave and differentiate according to their locations. Evidence going back over a century suggests that positional information is often specified in relative, rather than absolute coordinates, i.e. scaled to the size of the territory being patterned (Cooke, 1981; De Robertis, 2006; Driesch, 1891; Inomata, 2017; Ishimatsu et al., 2018; Teleman and Cohen, 2000). Fifty years ago, Wolpert argued that this implies that morphogen gradients adjust themselves to fit the fields on which they act (Wolpert, 1969). More recently such “morphogen gradient scaling” has been directly observed (Ben-Zvi et al., 2011b; Gregor et al., 2005; Hamaratoglu et al., 2011; Teleman and Cohen, 2000; Wartlick et al., 2011).

Wolpert noted that one type of gradient—a linear diffusion gradient from a source to an absorbing sink—scales naturally, automatically readjusting its slope when the location of the sink is moved (Wolpert, 1969). Over time, the concept that morphogens form linear source-to-sink gradients gave way to the current view—supported by observations (e.g. Eldar et al., 2002; Entchev et al., 2000; Gregor et al., 2007; Teleman and Cohen, 2000)—that gradients are shaped by continual decay throughout the morphogen field. Gradients shaped in this manner should not scale automatically, implying that some additional mechanisms exist to make scaling happen. Progress toward identifying possible mechanisms has been slow, but received a boost with the development of the Expansion-Repression (ER) model (Ben-Zvi and Barkai, 2010). In this model, a morphogen represses the expression of a secreted “expander”, which diffuses back toward the morphogen source, promoting morphogen spread (either by increasing morphogen diffusivity, or inhibiting morphogen uptake). This mechanism implements homeostatic feedback (moving a distal field boundary away from a morphogen source increases production of the expander, which spreads the morphogen gradient toward the field boundary). Moreover, if the expander is a long-lived substance, this mechanism approximates integral negative feedback control, a strategy that achieves “perfect” scaling (Ben-Zvi and Barkai, 2010).

So far, the ER model has been used to explain how gradients of morphogens of the bone morphogenetic protein (BMP) family scale in response to embryo manipulations (Ben-Zvi et al., 2008) and the normal growth of larval wing imaginal discs (Ben-Zvi et al., 2011a). In the latter case, the relevant morphogen is decapentaplegic (Dpp), which is produced by a stripe of cells in the center of the disc, spreading bidirectionally through the columnar epithelium to create gradients in the anterior and posterior compartments. During the larval period, wing discs grow at least 60-fold in anteroposterior dimension, and the Dpp gradient appears to scale along with it (Hamaratoglu et al., 2011; Wartlick et al., 2011). Support for the ER model in this setting was provided by the identification of a putative expander, the secreted protein Pentagone (Pent, also known as Magu). *Pent* expression appears to be directly repressed by Dpp, restricting *Pent* to the most lateral cells of the wing pouch (the part of the disc patterned by Dpp). Loss of *pent* leads to a dramatic shortening of the Dpp signaling gradient, and overexpression of *pent* expands the gradient (Vuilleumier et al., 2010). Studies by two groups have shown that, in the absence of *pent*, developmental scaling of the Dpp gradient is greatly impaired (Ben-Zvi et al., 2011a; Hamaratoglu et al., 2011). Although Pent’s mechanism of action is not fully understood, it has been shown to bind heparan sulfate proteoglycans (HSPGs) and trigger their removal from the cell surface. Such proteoglycans act as co-receptors for BMPs (Kuo et al., 2010), including Dpp (Fujise et al., 2001; Jackson et al., 1997), strongly suggesting that Pent expands Dpp gradients by inhibiting receptor-mediated Dpp uptake.

Here we re-examine the role of Pent in the *Drosophila* wing disc, focusing on an important requirement of the ER model, which is that the expander spread in an essentially uniform manner across the morphogen field. Spreading uniformly is not the same as merely being diffusible, as spread quantifies the balance between transport (e.g. diffusion) and decay, where decay refers to all processes that eliminate a substance from a diffusing pool (i.e. destruction, uptake, leakage out of the system). In a stable diffusion gradient, a common measure of spread is the “apparent decay length”, or *λ*_app_, the distance over which concentration falls by a factor of 1/*e* (Lander et al., 2009). In the ER model, if the expander’s *λ*_app_ is not greater than the size of the morphogen field, the expander will affect the morphogen differently at different locations, leading to a distortion of morphogen gradient shape and, if *λ*_app_ is small enough, ineffective scaling.

As described below, we find *λ*_app_ for Pent to be very small, strongly suggesting that the mechanism responsible for scaling the Dpp gradient cannot rely on Pent fulfilling the expander function required by the ER model. After carrying out a variety of genetic experiments, and exploring mathematical models, we arrived at an alternative model, in which the feedback that drives scaling is not repression of an expander, but morphogen-mediated regulation of receptor (and co-receptor) function, a phenomenon that is fairly widely observed in patterning systems (e.g. Cadigan et al., 1998; Fujise et al., 2003; Kahkonen et al., 2018; Lecuit and Cohen, 1998). A key feature of this model is that it is dynamic, terminating scaling at a size that depends on the parameters of the system. In view of evidence that a growing Dpp gradient itself participates in driving disc growth (Wartlick et al., 2011), this feature suggests ways in which bi-directional coupling between patterning and growth can be achieved.

## RESULTS

### Quantifying morphogen gradient scaling

Scaling can be quantified in various ways. For discrete pattern elements, such as gene expression boundaries, one can define it as the preservation of the relative positions of such elements. During developmental growth, however, sharp gene expression boundaries may only emerge late (del Alamo Rodriguez et al., 2004; Oliveira et al., 2014), or read out morphogen signals in indirect (e.g. time-integrated) ways (e.g. Balaskas et al., 2012; Dessaud et al., 2007; Nahmad and Stathopoulos, 2009). To investigate developmental scaling directly, it is therefore important to monitor morphogen gradients themselves, or gradients of immediate downstream signals (e.g. phosphorylated Mad [pMad], in the case of Dpp). As smooth gradients lack landmarks with which to assess relative position, doing so ideally means tracking the locations of constant gradient amplitudes over time. It can be challenging, however, to measure absolute concentration accurately in complex tissues. Moreover, absolute morphogen or signaling molecule concentration may not even be the best read-out of positional information across developmental time, because the cells that decode that information can change dramatically over the period of observation. This is especially true in wing discs, where individual cells grow at least 7 fold in volume between late first larval instar and the end of disc growth (Widmann and Dahmann, 2009).

For these reasons, morphogen gradient scaling is often quantified by measuring relative, rather than absolute, change in gradient shape. Typically, scaling is judged by the degree to which the apparent decay length (*λ*_app_) of the morphogen (or its downstream signaling intermediates) changes proportionately with size of the morphogen field (e.g. Wartlick et al., 2011). For gradients that are exponential in shape, *λ*_app_ corresponds to the constant *λ* in the formula *C*=*C*_*0*_*e*^-*x/λ*^ where *C* is concentration, *C*_*0*_ is a constant, and *x* is distance from the morphogen source. In practice, *λ*_app_ is often measured as the distance over which the gradient falls to 1/*e* of its starting value (or by extracting the decay length constant from a best-fit exponential curve). Here we follow others in using *λ*_app_ as a first-line metric for assessing scaling, but also take care to visualize and analyze absolute gradient shapes whenever possible (and especially when analyzing mathematical models). As we argue below, changes in gradient amplitude and shape may play an important role in enabling certain kinds of scaling mechanisms.

### Evaluating the role of Pentagone as an “expander” in morphogen gradient scaling

The ER model of morphogen gradient scaling requires an expander that spreads uniformly across the morphogen field, i.e. traverses the gradient without much decrement in concentration. Other studies have shown that Pent can be detected at a distance from its site of synthesis (Norman et al., 2016; Vuilleumier et al., 2010), and that overexpression of Pent in the posterior half of the disc can influence gradient shapes in the adjacent anterior half (Vuilleumier et al., 2010), but neither observation actually speaks to how Pent concentrations decline over distance.

To address this, we first selectively knocked down *pent* in the posterior compartment of the disc. We reasoned that, if Pent truly diffuses freely across the disc, then both sides of the disc must be fed by both the anterior and posterior sources of the molecule. Thus, the extent to which strong phenotypes from knocking down Pent in one compartment were observed in both compartments would provide a measure of how uniformly Pent spreads.

We used posterior compartment-specific drivers to express two *pent*-directed RNAis exclusively in the posterior compartment of wing discs and, as a positive control, a disc-wide driver to express RNAi everywhere. Effects on the Dpp signaling gradient were scored as changes to *λ*_app_ of pMad. The results (Figure 1) show that posterior knockdown of *pent* produces exclusively posterior effects—equivalent to those of global knockdown—and no detectable anterior effects. This implies that Pent decay is sufficiently strong, over distance, that at best a small fraction of Pent produced in the posterior reaches the anterior compartment. The alternative explanation that there is some barrier to Pent diffusion at the midline can be dismissed given the ease with which over-expressed Pent in the (entire) posterior compartment can produce phenotypes in the anterior (Vuilleumier et al., 2010).

**Figure 1:**
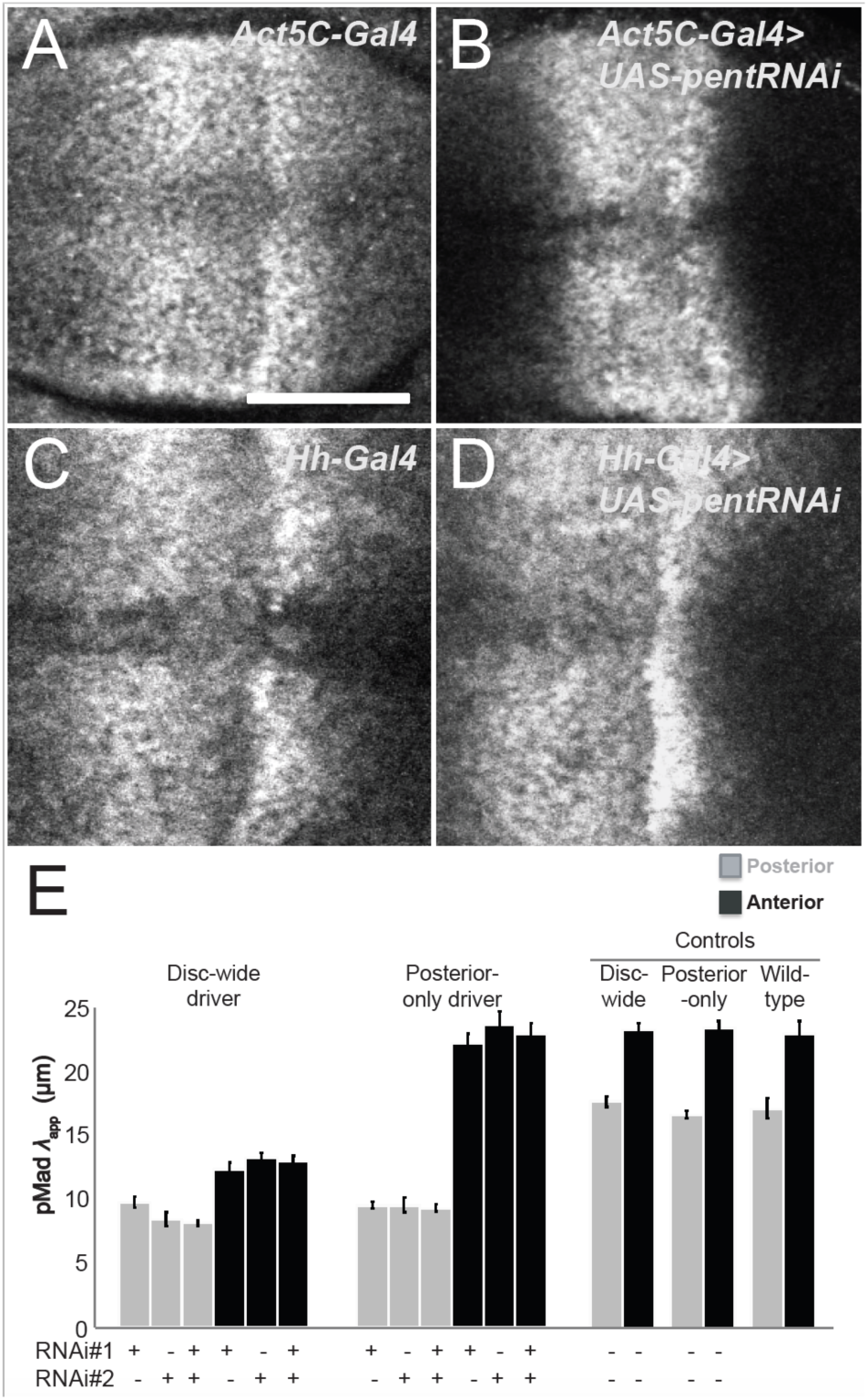
The effect of Pent on the Dpp signaling gradient is compartment-specific. (**A-D**) Phospho-Mad (pMad) staining of *Act5C-Gal4* (A), *Act5C-Gal4*>*UAS- pentRNAi* (B), *hh-Gal4* (C), and *hh-Gal4*>*UAS-pentRNAi* (D) third instar wing discs. Expressing *pentRNAi* (either of two different RNAi sequences) disc-wide (with *Act5C- Gal4*), expands the pMad gradient in both anterior and posterior compartments, whereas limiting *pentRNAi* to the posterior compartment (with *hh-Gal4*) affects only the posterior compartment. Bar = 50 µm. (**E**) Results are summarized as changes in pMad apparent decay length (*λ*_app_) in each compartment for each of the four genotypes in A-D. *N* varies between 12 and 23 for each condition. Error bars represent SEM.

If Pent concentrations decay over distance, then Pent itself should form gradients. To visualize these, we took two approaches. We first used a green fluorescent protein-tagged version of Pent (GFP-Pent), which we verified complements a *pent* mutation [Fig. S1]. We expressed GFP-Pent at various locations in wing discs using a variety of *Gal4* drivers (Figures 2A-C’ and Fig. S2A-D). From fluorescence intensity measurements acquired by confocal imaging, values of *λ*_app_ for GFP-Pent were obtained by fitting to a declining exponential function. In all cases, we observed a *λ*_app_ for Pent in the range of 6-8 µm. (Figure 2D and S2E).

**Figure 2:**
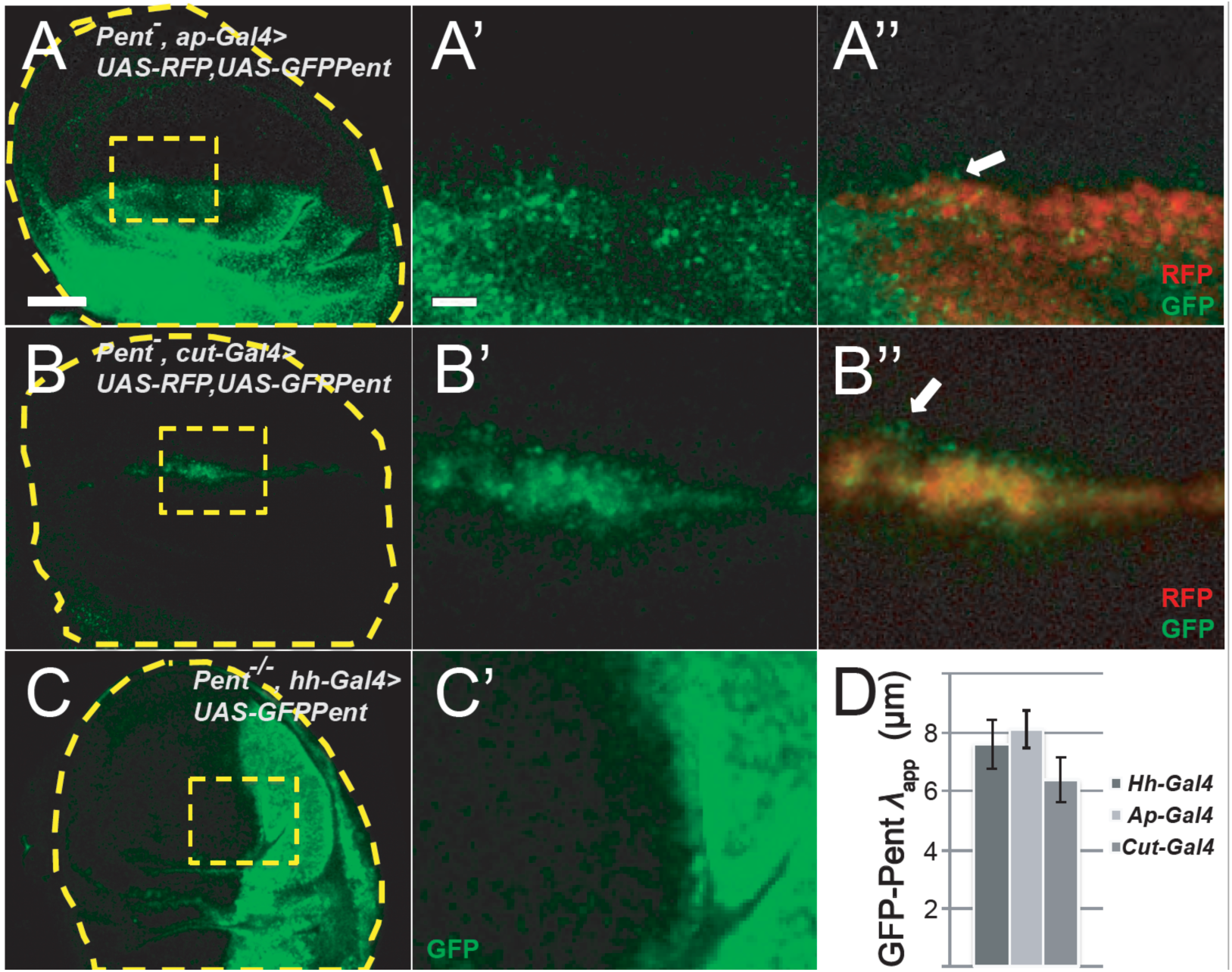
The apparent decay length (*λ*_app_) of GFP-Pent is relatively short. (**A-C**) *UAS-GFP-Pent* was expressed in various sharp patterns using specific *Gal4*-drivers: *ap-Gal4* (dorsal domain, A-A’’); *cut-Gal4* (dorsal-ventral boundary, B-B’’); and *hh-Gal4* (posterior compartment, C-C’). With *ap-Gal4* and *cut-Gal4, UAS-RFP* was used to mark the GFP- Pent-producing cells (A’’ and B’’, respectively). Arrows show GFP fluorescence located a few cell diameters away from the RFP-expressing cells. With *hh-Gal4*, both endogenous *pent* alleles were mutant, to show that the spread of GFP-Pent is similar whether or not all or only some Pent molecules were GFP-labeled. (**D**) Values of *λ*_app_ for GFP- Pent were determined for the genotypes in A-C, and were found to be between 6 and 8 µm. Bar in (A) = 50 µm and applies to panels A, B and C. The bar in A’ = 10 µm and applies to A’, A’’, B’, B’’ and C’.

Since it is possible that the diffusion or decay of Pent could have been affected by its fusion to GFP, we carried out further experiments that did not require tagging the molecule: we expressed wildtype *pent* in mosaic clones in wing discs. As shown in Fig. 3, *pent* overexpression clones are associated with two phenomena: reduced pMad staining (Figure 3A-E’), and reduced immunostaining for the cell-surface HSPG Dally-like protein (Dlp), a (Figure 3F-3I’). As discussed above, Pent binds to (Vuilleumier et al., 2010) and drives the internalization of both Dlp and the related HSPG Dally (Norman et al., 2016), so the observed loss of Dlp in response to Pent expression agrees with previous studies. The reduction in pMad staining, although not previously reported, is likely a consequence of the same process, since Dally and Dlp act as co-receptors for Dpp signaling, and their elimination in clones has previously been shown to produce cell-autonomous decreases in pMad (Belenkaya et al., 2004; Fujise et al., 2003).

**Figure 3:**
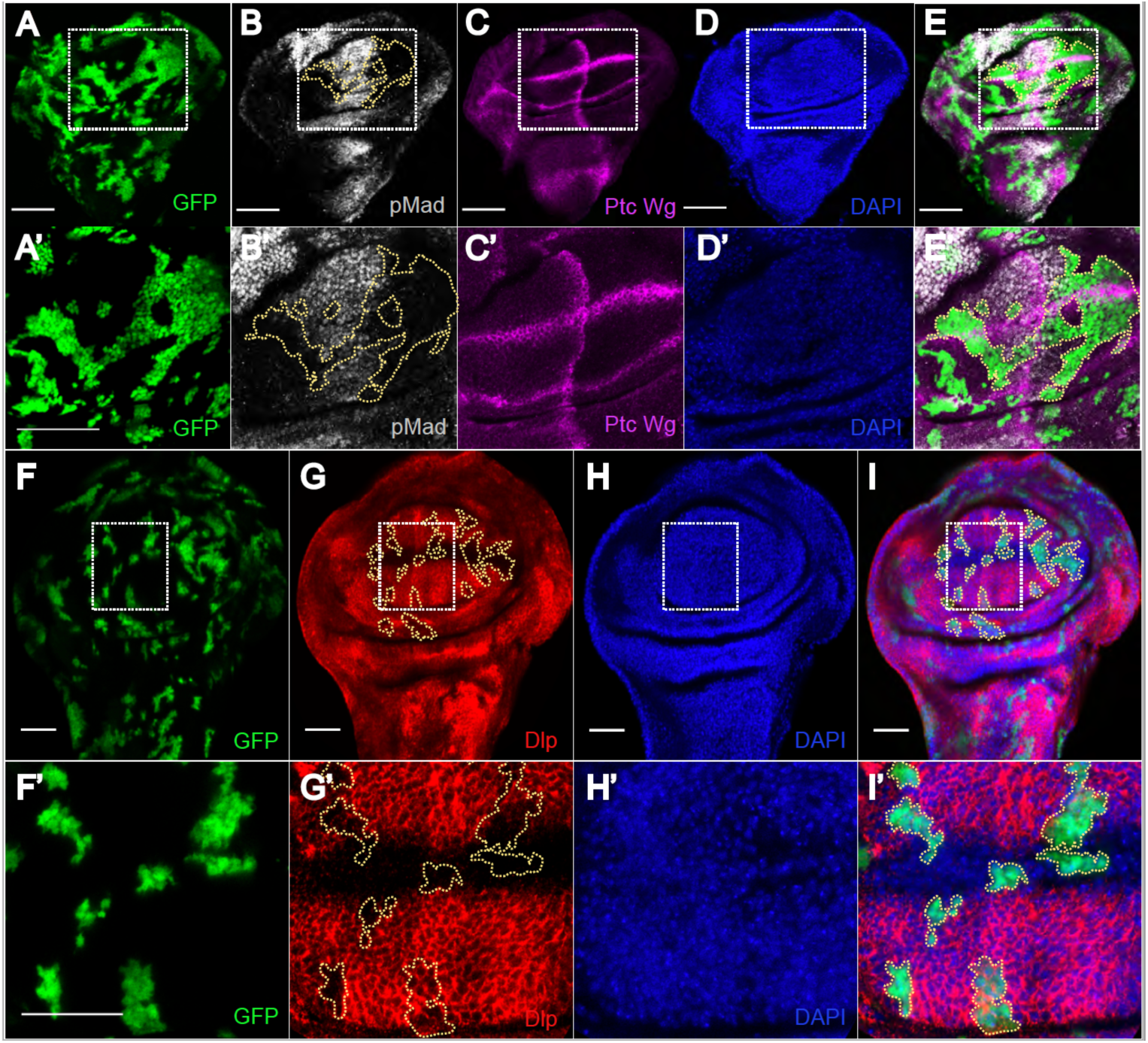
Pent overexpression has only local effects. (**A-E’**) Third instar wing disc with clones of cells overexpressing *pent* (GFP, green) stained with anti-pMad (gray), anti-Ptc and anti-Wg antibodies (magenta), and DAPI (blue). Ptc/Wg staining marks the A/P and D/V boundaries, as well as the Dpp production region. The rectangular region in panels (A-E) is blown up in panels (A’-E’). pMad staining can be seen to be markedly reduced within the clones (outlined area in B and B’) but only barely outside their boundaries. (**F-I’**) A wing disc containing clones of cells overexpressing *pent* (GFP, green) stained with anti-Dlp antibody, and DAPI (red and blue). The rectangle region in panels (F-I) is blown up in panels (F’-I’). Dlp-staining is markedly reduced within the clones (outlined areas in G and G’) but, again, almost no effect is observed outside of clone boundaries. Bars = 50 µm.

Close inspection of the clones in Fig. 3 shows that the effect of *pent* overexpression on pMad and *dlp* is nearly cell-autonomous, despite the fact that Pent is diffusible. Although a precise decay length for Pent cannot be derived from such experiments, the results suggest it is at most a few cell diameters (2-4 µm), if anything even smaller than *λ*_app_ estimated using GFP-Pent (Fig. 2). We conclude that, although Pent is diffusible, it decays strongly with distance. This suggests Pent acts relatively locally, rather than globally, undermining support for the ER model as an explanation for Dpp gradient scaling.

### Scaling of the Dpp signaling gradient is a transient phenomenon

Before investigating alternative explanations for gradient scaling, we more closely examined scaling dynamics. We collected a large number of wild type wing discs, spanning a range of sizes, and calculated *λ*_app_ for pMad, using the same measurement approaches taken by other investigators, making an effort to avoid various pitfalls and artifacts (e.g. measuring too close the dorsoventral boundary (Hamaratoglu et al., 2011)).

Our observations (Fig. S3; see also Fig. 4-5) confirm that *λ*_app_ grows roughly in proportion to disc size, but also indicate that it does so only up to the time (part-way through third larval instar) that posterior compartment sizes reach about 50-60 µm—about a fourth to a third of their final size, or about two cell cycles (approximately one day) prior to the end of disc growth. After that, scaling seems to cease rather abruptly. Such behavior has not been noted previously, possibly because other groups have focused more on documenting the existence of scaling during most of disc growth, rather than on how it behaves during the last stages of growth. What this behavior suggests is that whatever mechanism accounts for scaling of the Dpp gradient, it is size-limited, i.e. has a maximal distance over which it functions. Thus, our challenge in explaining scaling in the wing disc involves not only understanding how it happens, but also why it stops when it does.

**Figure 4:**
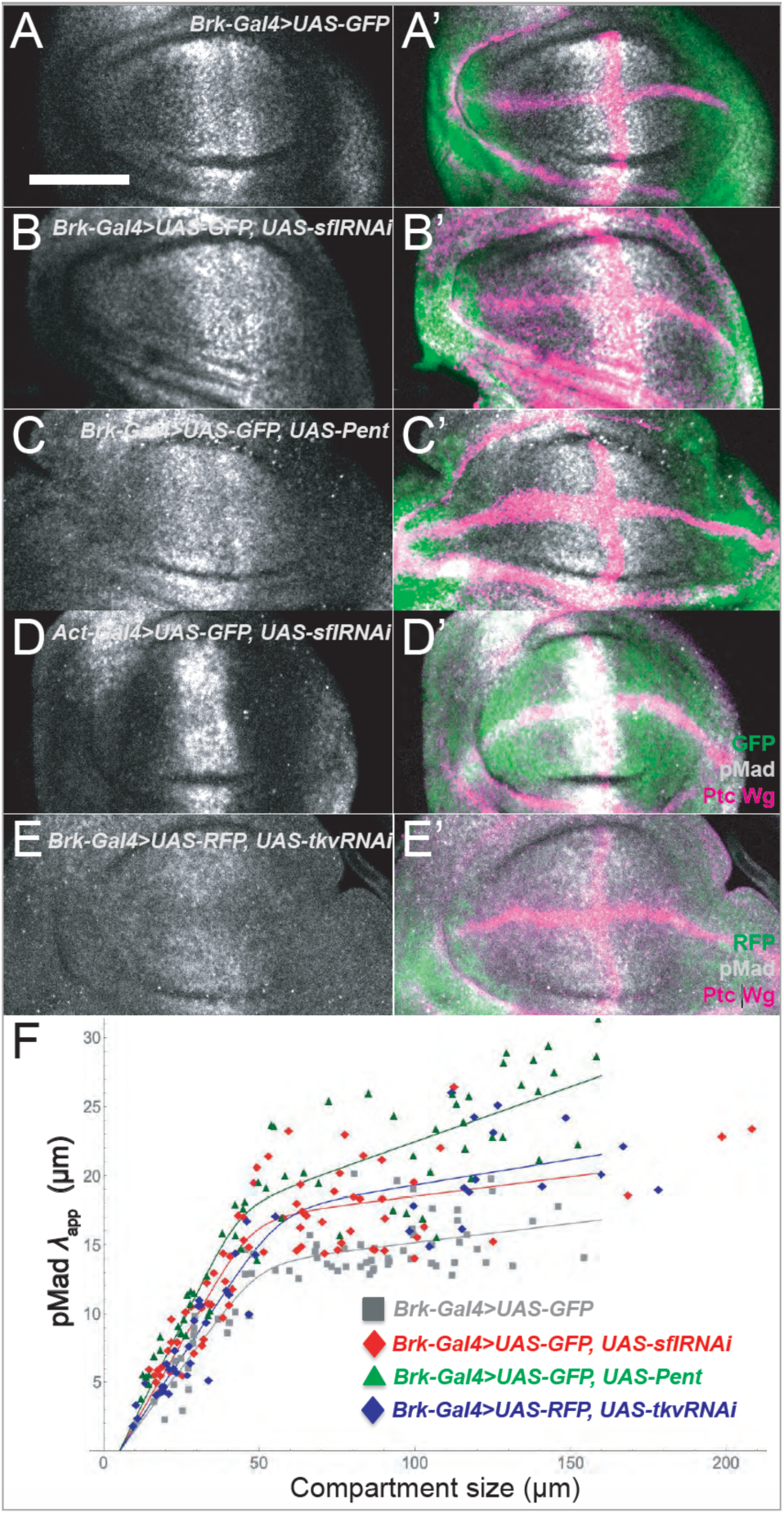
Local, cell-autonomous inhibition of co-receptor or receptor function phenocopies *pent* overexpression. (**A-E’**) *brk-*Gal4, which drives expression in a domain similar to that of *pent*, was used to drive expression in the wing disc of *UAS-GFP* (negative control, A-A’); *UAS-GFP; UAS-sflRNAi* (inhibition of HSPG sulfation B-B’), *UAS-GFP; UAS-pent* (*pent* overexpression, C-C’), and *UAS-RFP; UAS-tkvRNAi* (inhibition of receptor expression, E-E’). In addition, *Act5C-Gal4* was used to drive *UAS-GFP; UAS- sflRNAi* throughout the disc (D-D’). (**F**) *λ*_app_ for pMad in the posterior compartments of wing discs of the genotypes in A, B, C and E (gray squares, red diamonds, green triangles and blue diamonds, respectively) were determined for a large number of discs of different sizes, and plotted against compartment size. Data are fitted to smooth curves as described in methods. Scale bar is 50 µm.

### Pent function can be mimicked by cell-autonomously disabling co-receptors at the edges of the Dpp morphogen field

The evidence that Pent does not spread far (Fig. 1-3) raises the question of whether, to carry out its function, Pent needs to spread at all. One way to address this might be to physically tether Pent so that it cannot move. A simpler approach is to test whether Pent function can be mimicked by replicating the actions of Pent in a purely cell-autonomous way in the domain in which Pent is produced. As discussed above, current consensus is that Pent acts by reducing the expression (and thereby the function) of HSPG co-receptors, in so doing decreasing Dpp uptake and increasing Dpp spread. We gathered a number of new, ancillary observations consistent with that view: For example, *pent* mutants display shrinkage of the Dpp gradient itself (not just the downstream pMad gradient; Fig. S4A-B); there is no change in Dpp diffusivity (as measured by fluorescence correlation spectroscopy) in *pent* mutant discs (fig. S4C-F); and there appears to be no significant direct interaction between Pent and the Dpp receptor Tkv (as quantified using cross-correlation raster scanning intensity correlation microscopy, a sensitive test for molecular co-diffusion; Fig. S4G-I).

To reduce HSPG function in a cell-autonomous way, we used *sulfateless-RNAi* (*sfl-RNAi*). *Sfl* encodes the enzyme required for the N-sulfation, and consequently the function, of HSPGs (Baeg et al., 2001; Lin and Perrimon, 1999). We compared pMad gradients in the posterior compartment of wing discs in which either *sfl-*RNAi, *tkv*-RNAi, *pent*, or GFP were expressed under the control of *brk- Gal4*, which drives expression in a domain very similar to that of *pent* itself (Figures 4A-E). A smaller set of experiments were also carried out using *ds-Gal4*, which drives expression in a similar domain, and yielded similar results (Fig. S5A-E). We collected data from a large number of discs (>60 per genotype) with posterior compartment sizes varying over a wide range, and plotted pMad decay lengths against compartment size (Figure 4F). To account for changes in scaling behavior over time, we fit the data to a function that switches smoothly from one linear slope to another (both the slopes and the switching point were fit to the data).

As expected, the value of *λ*_app_ for *brk*-*Gal4*>UAS-*Pent* gradients was larger, at comparable disc sizes, than for control discs (*brk*-*Gal4*>UAS-GFP), showing that adding Pent in its own expression domain can further expand the Dpp gradient. Interestingly, *brk*-*Gal4*>UAS-*sflRNAi* gradients were similarly expanded, although not by quite as much as *brk*-*Gal4*>UAS-*Pent* gradients. This was true in small discs undergoing scaling (posterior compartment sizes from 10-50 µm) as well as large discs in which scaling had already slowed or stopped. We also noticed that adult wing phenotypes of *brk Gal4*>UAS-*sflRNAi* flies resembled those of *brk Gal4*>UAS-*Pent* flies (Fig. S5F-I).

These results argue that impairing HSPG function in the domain where Pent is expressed can phenocopy Pent overexpression. Consistent with the view that the role of HSPGs is to enhance receptor-mediated Dpp uptake, we also find that knocking down receptor expression in the *brk* domain has the same effect on the pMad gradient as disrupting HSPG function in the same domain (Fig. 4E, F).

Interestingly, if we express *sflRNAi* throughout the entire disc (using *Act5C*-*Gal4*), we observe shrinkage of the Dpp gradient (Fig. 4D’). This is a useful result because it argues that the expansion we see with *brk*-*Gal4*>UAS-*sflRNAi* (Fig. 4) and *ds*-*Gal4*>UAS-*sflRNAi* (Fig. S5) could not have been an artifact of “leaky” expression of these drivers in the central part of the disc, as such leaky expression should have produced the opposite result to what was seen.

These experiments also argue against a possible model of Pent action where Pent-mediated internalization of HSPGs does not actually destroy these molecules, but causes them to be released from cells in shed forms that then diffuse away, potentially acting as secondary expanders of the Dpp gradient. This idea is not implausible, given that cells do shed HSPGs of the family that includes Dally and Dlp (Bernfield et al., 1999; Ishihara et al., 1987); shed forms have been proposed to play a role in morphogen gradient formation (Giraldez et al., 2002); and a truncated, soluble form of Dally, when expressed in wing discs, can both diffuse widely and expand Dpp gradients (Takeo et al., 2005). But if this model were correct, eliminating HSPG function in the periphery of the wing disc should have contracted, not expanded, Dpp gradients, as we observed.

### Feedback regulation of receptors and co-receptors is required for scaling

One reason the apparent decay length, *λ*_app_, is widely used as a measure of morphogen gradient shape is that a simple model, known as the uniform-decay model, connects this metric to the underlying biophysics of gradient formation. Specifically, if morphogen decay is uniform in space, and the morphogen field is sufficiently large, then the steady state shape of a gradient should have the form *e*^*-x/λ*^, with 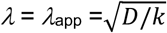, where *D* is the morphogen diffusion coefficient, and *k* is a rate constant of morphogen removal (e.g. uptake) (Lander et al., 2009).

In general, however, it is not valid to assume that morphogen decay is uniform in space, because morphogens can influence their own uptake or destruction. For example, some morphogens (e.g. Wingless and Hedgehog in *Drosophila* wing discs) upregulate their own removal; such “self-enhanced decay” characteristically distorts gradient shape (Eldar et al., 2003; Lander et al., 2009). In contrast, Dpp represses the synthesis of the Dpp receptor Tkv (through indirect effects of Dpp target genes on *tkv* transcription (Lecuit and Cohen, 1998)), as well as the synthesis of the HSPG Dally (Fujise et al., 2003). Tkv appears to be the major determinant of Dpp uptake and, thereby removal (Akiyama et al., 2008; Lecuit and Cohen, 1998), and co-receptors such as Dally can be expected to boost this function of Tkv, or even mediate some uptake themselves (Fujise et al., 2003). Consequently, we expect Dpp gradients to be distorted as the result of “self-repressed decay”.

The shape changes caused by self-enhanced and self-repressed decay may seem subtle when viewed in static graphs (Lander et al., 2009), but they can have a large effect on how gradients respond to perturbations. For example, self-enhanced-decay gradients display increased robustness to changes in amplitude (i.e. threshold locations do not move nearly as much as they do in uniform decay gradients; (Eldar et al., 2003)). Self-repressed-decay gradients, it turns out, can be expected to display enhanced sensitivity to changes in the size of the morphogen field, a phenomenon that—as we will see shortly—can drive morphogen gradient scaling.

Before discussing the theoretical grounds for this assertion, we present experimental evidence in support of it: To test whether self-repression of decay in the Dpp gradient of the wing disc is required for gradient scaling, it was necessary to disable the two feedback loops that lead from Dpp signaling to down-regulation of the expression of *tkv* and *dally*. For *tkv*, we took advantage of an existing transgenic allele that uses a Ubiquitin promoter to drive ubiquitous, unregulated expression of HA-tagged *tkv* (Ogiso et al., 2011). When combined with homozygous null mutation of the endogenous *tkv* locus, viable flies are obtained, with late-third larval instar wing discs that do not differ significantly in pattern from wildtype, except for the fact that *tkv* expression is spatially uniform, rather than graded. We refer to this genotype as “Ubi-*tkv*”.

To disable feedback on *dally*, we obtained a UAS-*dally* transgene, and used an *Act5C-Gal4* driver to drive its expression in a uniform pattern in a *dally*-mutant background (*dally*^*80*^*/dally*^*80*^). We refer to this genotype as “Uniform-*dally*”.

Figure 5 shows the results obtained from evaluating a large number of wildtype, *pent, pent*^+/-^, Ubi-*tkv*, Uniform-*dally*; Ubi-*tkv*/Uniform-*dally*; and *pent*/Ubi-*tkv* and *pent*/Ubi-*tkv/*Uniform-*dally* discs of a broad range of sizes. *λ*_app_ was measured for pMad gradients and plotted against posterior compartment sizes. Curves were fit as in Fig 4I and 5A, allowing the size at which scaling behavior slows or stops to be estimated independently for each data set.

**Figure 5:**
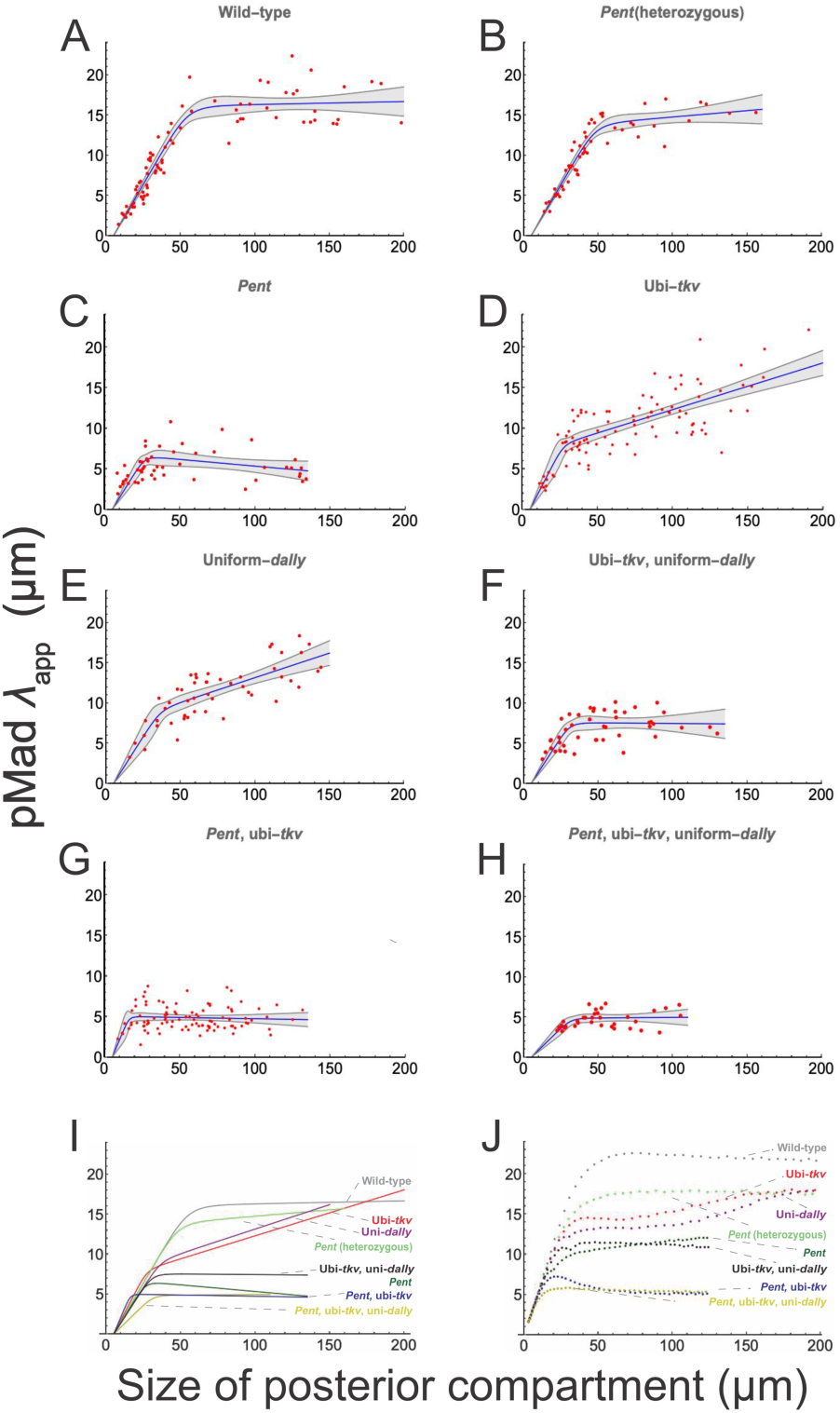
Feedback regulation of *tkv* and *dally* is required for scaling of the pMad gradient. (**A-H**) pMad decay lengths in the posterior compartment were measured for a large number of wing discs at different developmental stages, for 8 genotypes: Wild-type (A); heterozygous *pent* mutant (B, *+/pent*^*2*^), homozygous *pent* mutant (C, *pent*^*2*^*/pent*^*2*^); ubiquitously-expressed *tkv* (D, *tkv*^*Df*^*/tkv*^*strII*^; *ubi-tkv-HA*); uniformly-expressed dally (E, *Act5C-Gal4, dally*^*80*^*/UAS-Dally, dally*^*80*^); ubiquitously-expressed *tkv* and uniformly-expressed *dally* (F, *tkv*^*strII*^*/tkv*^*a12*^, *ubi-tkv- HA; Act5C-Gal4, dally*^*80*^*/UAS-Dally, dally*^*80*^); ubiquitously-expressed *tkv*, homozygous pent mutant (G, *pent*^*2*^, *tkv*^*Df*^*/pent*^*A17*^, *tkv*^*strII*^; *ubi-tkv-HA*); ubiquitously-expressed *tkv*, uniformly-expressed *dally*, and homozygous pent mutant (H, *pent*^*A17*^,*tkv*^*strII*^*/pent*^*2*^, *tkv*^*a12*^, *ubi-tkv-HA; Act5C-Gal4, dally*^*80*^*/UAS-Dally, dally*^*80*^); Data are fitted to smooth curves, as in Fig. 4. Envelopes around curves represent 95% confidence intervals for the fits. (**I**) Summary of results in panels A-H. (**J**) Data obtained from mathematical modeling (see Supplementary Data, and Fig. 6), showing the evolution of the pMad apparent decay length over time.

The results indicate that scaling is significantly impaired when either *tkv* or *dally* regulation is bypassed, and nearly eliminated when both are bypassed together. These differences emerge mainly after posterior compartments have exceeded 30 µm in size. Above that size, Ubi-*tkv* and Uniform-*dally* continue to scale, but much more slowly than wildtype gradients. Eventually, however, such gradients do “catch up” to wildtype gradients, as a result of the fact that wildtype gradients cease scaling sooner. In contrast, doubly-mutant Ubi-*tkv*/Uniform-*dally* gradients stop expanding altogether once posterior compartments grow beyond about 30 µm, reaching a final *λ*_app_ about half that of wildtype.

The defect in Ubi-*tkv*/Uniform-*dally* gradients is almost, but not quite, as severe as that in *pent* mutants, which cease scaling at a slightly earlier stage. Interestingly, the phenotype of *pent*/Ubi-*tkv* and *pent*/Ubi-*tkv/*Uniform-*dally* discs was only slightly more severe than for *pent* alone. These results strongly support the conclusion that both scaling, and the effect of Pent on scaling, depend upon feedback regulation of *tkv* and *dally* by Dpp.

### Modeling the dynamics and endpoints of scaling

To explain the behaviors shown in Fig. 5, we turned to mathematical modeling. Accounting for all the cell biological phenomena that affect Dpp gradient shape requires modeling a rather large number of molecular species and processes. As many of these processes are not quantitatively well understood, they were represented as simply as possible, and model behaviors were explored over parameter ranges that were wide but plausible (given the available data). The goal of such modeling is not so much to identify parameter values as to determine whether existing observations can be matched without invoking additional mechanisms. To the extent that inclusion of new mechanisms is not required, models such as these can help us identify which processes potentially play the most important roles in morphogen gradient scaling.

The molecular species represented in the model are Dpp, Tkv, “co-receptor” (to represent both Dally and Dlp), Pent, and pMad, Dpp-coreceptor complexes, and two types of Dpp-receptor complexes (the more stable of which forms with the aid of coreceptor-mediated catalysis; (Kuo et al., 2010)), plus pMad and Brk. An additional transcription factor is included downstream of Brk to enable Brk (which is a repressor) to indirectly activate Tkv and co-receptor synthesis (we do not name this factor explicitly, but its role in the model resembles that of *optomotor blind* (del Alamo Rodriguez et al., 2004)). Only Dpp is modeled as a diffusing species, as Pent’s own decay length is so short. Details of the model are explained in the supplemental material.

Summary results for multiple genotypes are shown in Figure 5J, with a detailed simulation shown in Fig. 6 (for parameter selection, see Supplemental Materials). The behaviors observed for each of the genotypes in Figure 5I are reasonably well replicated: Initially, all modeled genotypes scale well, until posterior compartment sizes reach ∼10 µm. Up to this time, Dpp and pMad gradient shapes produced by the model are essentially straight lines from source to the end of the morphogen field (Fig 6). Automatically-adjusting straight line gradients call to mind the “source-sink” scaling mechanism of (Wolpert, 1969). Such scaling exemplifies what mathematicians call a “boundary layer effect”, whereby phenomena at a boundary influence gradient shape at a distance. For steady state diffusion gradients, the approximate distance over which boundary layer effects occur is given by the *intrinsic decay length constant, λ*_intrinsic_, defined as the square root of the ratio of the diffusion coefficient and the (effective) decay rate constant.

**Figure 6:**
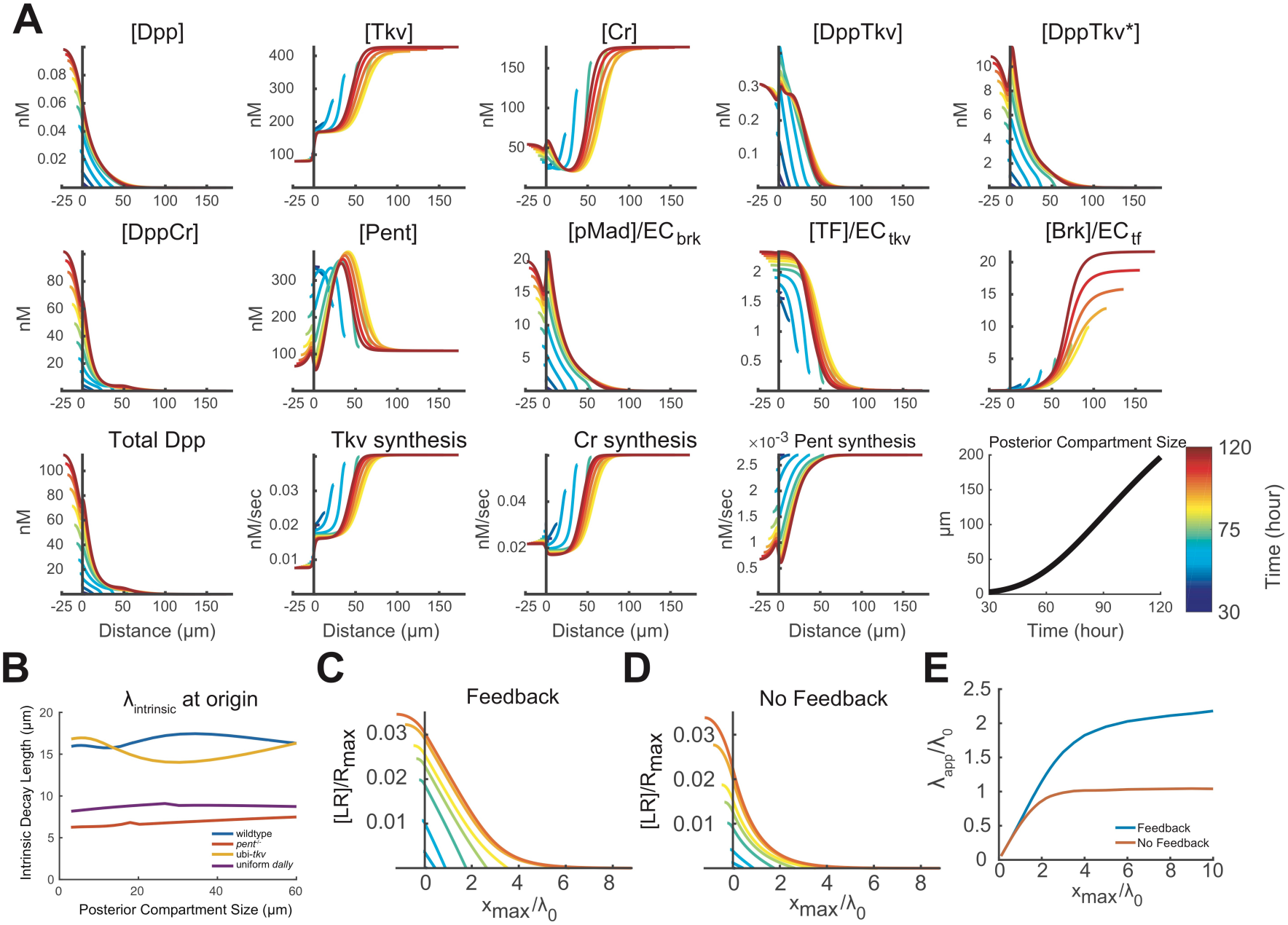
Modeling Dpp gradient scaling in the posterior compartment of the wing disc. **A**. Time evolution of the wildtype Dpp morphogen gradient. The graphs show the distributions of free Dpp, Dpp-receptor and -coreceptor complexes, downstream signals and targets, simulated according to the model given by equations (1) in Supplemental materials. *x*=0 represents the border between the Dpp production region and the posterior compartment. The final graph in the simulation shows the growth of the posterior compartment over time, and the legend shows how time is represented as color in each of the graphs. Notice how Dpp and pMad gradients are relatively linear at early times, but later become more exponential in shape. Simulations for other genotypes are shown in the Supplemental material. **B**. The Dpp intrinsic decay length (*λ*_intrinsic_) for four genotypes (wildtype, *pent*^*-/-*^, ubi-*tkv* and uniform-*dally*), was calculated as a function of time, and plotted as a function of compartment size (*x*_max_). *λ*_intrinsic_ captures this distance over which boundary effects occur, so that *λ*_intrinsic_/*x*_max_ provides a measure of the extent to which a gradient’s shape is boundary-controlled. Note that, in wildtype, *λ*_intrinsic_ transiently rises, reflecting the effects of feedback downregulation of *tkv* and *dally*. **C-E**. Illustration of pseudo-source-sink scaling using a simplified model. These panels show the output of a much-reduced, steady-state version of the model that retains only ligands, receptors and ligand-receptor complexes; irreversible capture of ligands by receptors; and downregulation of receptor synthesis by ligand-receptor complexes. The reduced model has only four free parameters. Values of *LR* (which stands for ligand-receptor complexes) are normalized to *R*_max_ (the receptor concentration that would be obtained in the absence of ligand binding or feedback), and plotted against compartment size normalized to the intrinsic decay length that would be observed in the absence of ligand binding or feedback (*λ*_0_). The example in panel C illustrates the effect of feedback downregulation of receptor production; in panel D feedback is turned off and ligand production rate adjusted to produce a similar value of LR. Panel E summarizes the apparent decay lengths (*λ*_app_), relative to *λ*_0_, for the curves in C-D. Notice how, as a result of feedback, the *LR* gradient achieves a much longer period of scaling. For further explanation and extended parameter space exploration see Supplemental Materials.

As we previously noted, for uniform-decay gradients on a sufficiently large field, gradient shape should be described by *e*^*-x/λ*^, with 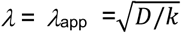; thus, for such gradients *λ*_app_ = *λ*_intrinsic_. But “sufficiently large field” here turns out to mean large compared with *λ*_intrinsic_. With fields smaller than *λ*_intrinsic_, gradient shape becomes less exponential, and more linear. Thus, the more linear the gradient, the farther into it boundary-layer effects will be observed. Wolpert’s source-sink gradients are merely the limiting case in which *λ*_intrinsic_ = ∞, (no decay within the morphogen field), yielding perfectly linear gradients that scale perfectly with movement of the boundary. As long as morphogen gradients operate in a regime of large *λ*_intrinsic_ (compared with morphogen field size), they too will scale automatically. However, this can only go on for so long, as field size should eventually catch up with *λ*_intrinsic_ –at which point gradients will start looking exponential and scaling will stop.

In the mathematical model, the scaling initially displayed by all genotypes stops at different sizes (in agreement with experimental observations; Fig. 5I), for reasons that depend on the genotype. We consider first the wild type: In that situation, the initial value of *λ*_intrinsic_ (∼16 µm everywhere) suggests that scaling should fail sooner than it does, but then the value of *λ*_intrinsic_ near the morphogen source rises as the disc grows (Fig. 6B), extending the period of scaling. The reason *λ*_intrinsic_ grows is that receptors and co-receptors become increasingly downregulated (since they are the primary contributors to morphogen decay, their loss drives *λ*_intrinsic_ up). Interestingly, to prolong scaling it is not necessary for *λ*_intrinsic_ to grow as fast as the disc itself. This is because once strongly non-uniform expression of receptors and co-receptors sets in—low near the morphogen source and high far away—the actual sink at the far end of the morphogen field becomes less and less important. Instead, the territory with high receptor/co-receptor expression itself acts like a sink, due to the higher level of morphogen uptake there. We call this behavior “pseudo-source-sink” scaling, as it emulates a boundary-layer effect without the need for a true tissue boundary (Fig. 6C-E).

The phenomenon fundamentally responsible for driving pseudo-source-sink scaling is amplitude growth: In other words, it is because Dpp and pMad levels at the start of the gradient rise with disc growth that receptor and co-receptor expression becomes repressed to a greater degree, and farther away, over time. Thus, whereas source-sink scaling exemplifies a direct coupling of field size to gradient scale, pseudo-source sink scaling depends on indirect feedback: changes in field size first produce changes in gradient amplitude, and these then drive changes in gradient scale.

Why should changes in field size cause changes in gradient amplitude? In the model, several processes contribute. The simplest is that the production region itself grows with the disc; as it does so, it feeds more Dpp into the gradient (the magnitude of the effect depending on the level morphogen decay within the production region). A second reason arises from the laws of physics and the fact that the gradient has already been scaling: According to Fick’s first law, net diffusive flux at any point is proportional to the slope of the diffusion gradient. So whenever a gradient expands by becoming shallower, the diffusive flux at the origin of the gradient must decrease. That in turn leaves more molecules available to contribute to the local concentration of morphogen, raising the free morphogen concentration.

Two other mechanisms can also contribute to amplitude growth, but are relatively minor contributors in the model: To the extent that Dpp molecules associated with or internalized within cells are very long-lived, the Dpp signal that cells receive will lag significantly behind the free Dpp concentration, which can cause pMad levels to rise even if Dpp levels are not. And to the extent that disc growth is not purely exponential, but rather slows as time goes on (Wartlick et al., 2011), the loss of Dpp and pMad due to dilution can be expected to go down, ultimately raising Dpp and pMad concentration.

Does amplitude growth, of the kind we observe in the model, actually happen in vivo? Directly comparing Dpp and pMad amplitudes at different time points is challenging, not only because of individual variation among discs, but because discs change dramatically in thickness as they grow, necessitating corrections for systematic changes in the efficiency of immunostaining and/or imaging. Nonetheless, groups that have produced such measurements consistently report amplitude growth in the wing disc Dpp gradient, although the degree to which they observe it varies (Hamaratoglu et al., 2011; Wartlick et al., 2011). Although the model parameters used in Fig. 5-6 predict an approximately 5-fold increase in Dpp and 10-fold in pMad over the corresponding time interval (Fig. 6), the actual changes are likely less important than the fold decrease they produce in Tkv and co-receptor expression. To get a sense of whether such a decrease actually occurs in vivo, we monitored beta-galactosidase expression in an enhancer trap line that reports on *tkv* expression, over a wide range of disc sizes. As shown in Fig. S7, the data support the assertion that, at early stages, *tkv* is not very strongly suppressed (and is consequently expressed throughout the disc), but it becomes so over time as discs grow. This both lowers *tkv* near the Dpp source, and continuously pushes the location of high *tkv* expression farther away from the Dpp source. These findings agree with those of (Widmann and Dahmann, 2009), who find that *brk* expression is also fairly uniform in early discs, and only becomes strongly suppressed by Dpp later.

Given that feedback regulation of receptors and co-receptors play an essential role in prolonging scaling in the mathematical model, it is not surprising that genotypes that eliminate both feedback loops stop scaling much earlier (at a posterior compartment size of ∼20 µm). In contrast, if only a single feedback loop is eliminated, gradients expand for a bit longer (posterior compartment size ∼30 µm), then very gradually catch up to a final *λ*_app_ almost equal to that of wild type (this agrees with experimental observations [Fig. 5]). Examination of the model explains this behavior: because a single feedback loop capable of adjusting *λ*_intrinsic_ remains, pseudo-source sink scaling remains possible, but the slower pace at which it happens means that one of the factors that contributes to amplitude growth (decreased diffusive flux due to shallower gradient slope) is less pronounced, leading to much slower scaling.

In the model, scaling also fails for the homozygous *pent* mutant, about when the posterior size reaches ∼20 µm, but the reason for this is entirely different. Because *pent* expression is repressed by Dpp, and Dpp signaling levels are low overall when discs are very small, *pent* is initially predicted to be expressed at high levels in most or all of the disc. Since Pent removes co-receptors, the absence of Pent increases co-receptor function, thereby decreasing *λ*_intrinsic_ in most or all of the disc, and causing source-sink scaling to fail prematurely. Thus, whereas elimination of feedback control of receptors and co-receptors impedes scaling by interfering with the process of scaling itself, elimination of Pent impedes scaling by changing the initial conditions of the disc. Consistent with this view are the results of RNAi up- and down-shift experiments (Fig. 7), which suggest that the effects of Pent on gradient scale are mainly due to actions early in disc growth (i.e. before mid-third instar).

**Figure 7:**
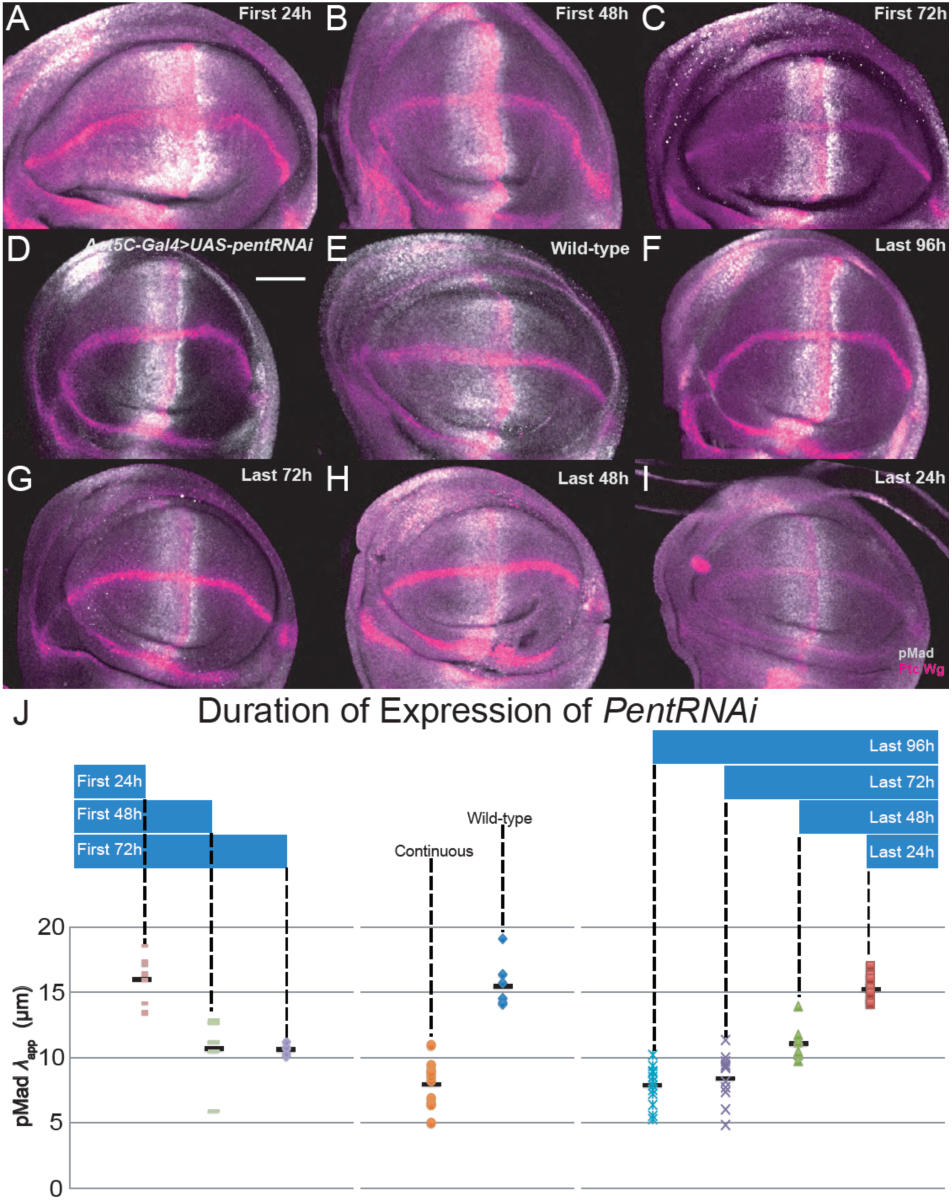
Investigating the temporal requirement for Pent. The *Act5C-Gal4; tubP-Gal80*^*ts*^ system was used together with *pentRNAi* to achieve inducible or repressible knockdown of *pent* (see Methods). (**A-I**) Representative wing discs in which *pentRNAi* was either expressed continuously (D), not expressed (E), or expressed for defined time periods—24 (A), 48 (B), or 72 hr after egg laying (C); or 96 (F), 72 (G), 48 (H) or 24 h (I)) prior to wandering stage. Discs were stained with anti-pMad (gray), and mixed anti-Ptc and anti-Wg antibodies (magenta). (**J**) Comparison of pMad decay lengths in the posterior compartments of wing discs from the above larvae. Bar = 50 µm.

In the model, the behaviors seen with all other genotypes may be understood as combinations of the effects discussed above. The one behavior observed in the experimental data that was not well reproduced in an earlier version of the model was the fact that once *pent* mutants stop scaling, they do not later “catch-up” in the way that Ubi-*tkv* and Uniform-*dally* genotypes do. Since feedback regulation of Tkv and co-receptors should be intact in *pent* mutants, we might have expected to see such behavior. However, when we included in the model the observations of (Hamaratoglu et al., 2011) that maximum expression of Brinker (*brk*) increases dramatically during disc growth, we saw that such catch-up behavior was suppressed. In effect, Brinker amplitude growth acts as a countervailing force to pseudo-source sink scaling, pushing thresholds back toward the Dpp source at the same time that increased Dpp signaling pushes thresholds farther away. As Hamaratoglu et al., point out, highest *brk* expression occurs where there is essentially no Dpp signaling, so the Dpp gradient cannot be the cause of the dramatic change in Brk amplitude over time. In the model, we arbitrarily adjust *brk* amplitude to follow the findings of (Hamaratoglu et al., 2011), however it is intriguing to speculate that there could be some coupling between *brk* expression and disc size that would make such coupling automatic.

## DISCUSSION

Morphogen gradients play a central role in animal development by tying cell behavior to spatial location. Whereas monotonic gradients of almost any sort can encode absolute location, for gradients to encode *relative* location on a domain of changeable size, gradients must scale to fit the territory they pattern, i.e. the morphogen field. For this to occur, at least one of the three processes that control gradient shape—morphogen production, transport or removal (decay)—must somehow be coupled to field size.

There have been several proposals for how this might occur automatically (Ben-Zvi and Barkai, 2010; Cheung et al., 2011; Wolpert, 1969), an elegant example of which is the expansion-repression (ER) model. The present study began as an attempt to test whether the secreted protein Pent, which has been implicated in scaling of the *Drosophila* wing disc Dpp gradient, exhibits the properties of the expander required by this model. We present experimental evidence that Pent lacks the necessary spatial range, and that Pent’s effects can even be phenocopied by disabling non-diffusible co-receptors in just the Pent expression domain [Figs. 1-4], suggesting that Pent need not diffuse to exert its influence on morphogen gradient scale. We go on to make the novel observation that feedback repression of receptor and co-receptor synthesis is required for scaling (Fig. 5].

How exactly, then, does scaling happen? Having reproduced the phenotypes of wildtype and mutant Dpp gradients with a mathematical model (Fig. 5-6) gives us the opportunity to develop a plausible explanation. What is most surprising about this explanation is that it does not end up attributing scaling to a single automatic mechanism, but rather to a collection of passive and active processes that collectively get the job done.

During the earliest phase of disc growth— corresponding approximately to first through early or mid-second larval instar–the model implements source-sink scaling because, when the disc is very small, *λ*_intrinsic_ is larger than the field size, allowing the distal boundary of the morphogen field to act as a sink, and gradient shape to become linear. How plausible is this scenario?

Although we cannot readily measure Dpp or pMad gradients in discs with posterior compartments smaller than 10 µm (at most a handful of cells), we do know that, from the earliest times at which measurements can be made, the shapes of the Dpp and pMad gradients that we can detect are not distinguishable from straight lines ((Hamaratoglu et al., 2011; Wartlick et al., 2011) and data presented here). And although we cannot confirm experimentally that scaling occurs prior to this point, we can say that, once discs reach this size, the pMad decay lengths of all genotypes are fairly similar (Fig. 5), which is consistent with source-sink scaling. Furthermore, we know both from published studies (Entchev et al., 2000; Hamaratoglu et al., 2011; Teleman and Cohen, 2000; Wartlick et al., 2011), and from the data collected here that, when discs reach their final sizes, *λ*_app_ for Dpp and pMad are on the order of 20 µm. As *λ*_app_ sets a lower bound on *λ*_intrinsic_, it is reasonable to think that in vivo parameters could easily produce *λ*_intrinsic_ that is larger than early field sizes (i.e. ≤10 µm).

As for the existence of a “sink” at the far end of the morphogen field, we think an effective sink is created by the geometry of the tissue. Assuming the primary space in which Dpp diffuses is the basolateral interstitial space (we have observed fluorescently-labeled Dpp in this space directly, and measured its diffusion there (Zhou et al., 2012)), then we know that that space is bounded apically by tight junctions (a “no-flux” boundary), but basally by a basement membrane through which molecular diffusion proceeds unimpeded. Dpp should thus constantly leak out of the basement membrane (Lander et al., 2011), a phenomenon that is supported by recent experimental measurements (Setiawan et al., 2018). Because the basement membrane wraps around the disc at its edges, when Dpp in this space reaches the anterior and posterior boundaries of the disc, it should also pass straight through into the hemolymph surrounding the disc, providing the functional equivalent of a sink.

At later times in the model, as discs grow beyond their initial values of *λ*_intrinsic_, wild-type discs transition to what we term “pseudo-source-sink scaling” (Fig. 6). Increasing levels of morphogen drive down receptor and co-receptor expression, increasing the value of *λ*_intrinsic_ near the morphogen source. Far away from the morphogen source, however, *λ*_intrinsic_ remains small, effectively creating a “pseudo-sink”. Gradients are still more-or-less linear, but no longer extend all the way to the disc boundary. As long as growth occurs in the amplitude of the morphogen gradient near the source, gradients respond by becoming shallower near the source, which makes them expand outward. Moreover, as discussed above, when gradients become shallower, that in itself drives amplitudes up, creating something of a positive feedback loop.

This process occurs until receptor and co-receptor expression can be downregulated no further (expression cannot be downregulated to zero because receptor and co-receptor function is required to drive the repression). After that, wildtype gradients outgrow their value of *λ*_intrinsic_, cease scaling altogether, and adopt a more-or-less exponential shape. Not surprisingly, genotypes that compromise the ability of the morphogen to repress receptor and/or co-receptor synthesis result in gradients that cease scaling earlier.

The idea that amplitude increases contribute substantially to morphogen gradient scaling has been proposed before, in connection with the Bicoid gradient of the *Drosophila* embryo. Selection for larger or smaller embryos is accompanied by partially compensatory shifts in the locations at which Bicoid target genes are turned on. These shifts occur not because of a change in Bicoid’s *λ*_app_, but rather an increase in gradient amplitude alone (Cheung et al., 2011).

For exponential gradients (like the Bicoid gradient), simple amplitude increase produces constant-distance shifts in threshold positions, in effect making thresholds near the morphogen source over-scale and those far away under-scale. Thus, like source-sink scaling, scaling due to pure-amplitude growth runs into spatial limits beyond which it is not very effective. The pseudo-source-sink mechanism partially compensates for this problem by displacing the effective sink away from the source as the amplitude near the source grows. However, it should be noted that in this case, what is truly scaling is the relative shape of the morphogen gradient, as measured by its *λ*_app_, and not the locations where absolute thresholds are crossed. At least some Dpp target genes (e.g. *dad*) do seem to scale in just this way (Hamaratoglu et al., 2011; Wartlick et al., 2011).

In addition to the various mechanisms— source-sink scaling, amplitude growth, pseudo-source-sink scaling—that drive Dpp gradient expansion in the model described here, additional processes may matter in vivo. Fried and Iber (Fried and Iber, 2014) argue that, because growth itself necessarily propels forward the molecules within a tissue—a process termed *advection—*growth-driven gradient expansion should occur by that process alone. Whether advective scaling is a large effect or—as in the model in Fig. 5J-6, a very small one—depends upon two time scales, which are determined by the parameters of the system: the relative growth rate of the field (i.e. tissue doubling time), and the relative “relaxation” rate of the morphogen or its downstream signals (how long it takes a local disturbance to decay away). The latter time scale is set by the effective half-life of the species in question. For diffusing molecules, relaxation is typically very fast, so that advection is not expected to have much influence on gradients of free morphogens, but for cell-associated species (receptor-bound morphogen, signaling intermediates), advective effects can be large if those species are sufficiently long-lived. There is a limit to how effective this phenomenon can be, however, because tissue growth necessarily causes dilution of all species, meaning that no species can have a half-life longer than the doubling time of the field itself.

Overall, the experiments and modeling presented here suggest a view of scaling as more “kluge” than elegant control system. Small fields with leaky boundaries can achieve source-sink and amplitude-growth scaling effectively “for free”, but not for very long, as both fail at large field size. Pseudo-source-sink effects can then kick in to prolong scaling.

The potential utility of source-sink-like behavior in morphogen patterning has been relatively neglected by modern developmental biologists, presumably because the exponential forms of late-stage morphogen gradients are better fit by a uniform decay model. Yet the present study suggests that the linear, source-sink gradients of early theorists (Wolpert, 1969) may sometimes be a better description of early stage gradients. Interestingly, recent work argues that source-sink behavior is also the primary determinant of BMP gradient shape in early zebrafish embryos (Zinski et al., 2017). The behavior in that system would, by our nomenclature, be more precisely termed pseudo-source sink, since the sink in that system arises from the binding of BMP to chordin, and chordin is downregulated by BMP—a feedback loop functionally analogous to the downregulation of Tkv and Dally by Dpp in the wing disc. The parallels between that system and the work described here are intriguing, because the BMP gradient that patterns the early vertebrate embryo also exhibits scaling behavior (in response to embryo bisection (Ben-Zvi et al., 2008; De Robertis, 2006)).

One of the drawbacks of using feedback downregulation of receptor function to extend the range of a morphogen gradient is that such feedback necessarily reduces the morphogen signal, which in turn lowers the feedback. Receptor function cannot therefore be lowered indefinitely, which places a limit on how large *λ*_intrinsic_ can be. Beyond this size, pseudo-source-sink scaling must fail. While this may seem like a deficiency, it can also be useful in the event that the morphogen gradient itself regulates disc growth, which for Dpp and the wing disc is well established (Affolter and Basler, 2007; Martin-Castellanos and Edgar, 2002). Indeed, Wartlick et al. argue that it is the continual rise in Dpp signaling within cells that is the primary signal that maintains growth (Wartlick et al., 2011). Because gradient scaling is required to ensure that this rise occurs proportionally at all locations, failure of scaling could potentially play a causal role in terminating growth. What is intriguing about this idea is that it predicts that the size at which growth stops should correlate with the size at which scaling stops, which we in fact observe in the data: As shown in Fig. 5, *Pent*-/- and Ubi- *tkv*/Uniform-*dally* mutant discs stop growing at a substantially smaller size than wildtype discs. These observations suggest that it may be more appropriate to view scaling and growth as a single coupled system, rather than a mechanism for adjusting pattern to match size.

Although the present study provides a potential explanation for Dpp gradient scaling that correctly predicts the scaling phenotypes of *pent* mutants, it does not provide a satisfying answer to the question of why Pent is utilized by wing discs in the first place. In the model, *pent* discs fail to scale because they start out with too small a value of *λ*_intrinsic_, but since all Pent does (in the model) is inhibit receptor function, discs should be able to obviate the need for Pent by just expressing a lower level of receptors or co-receptors. Why this isn’t what wing discs do is unclear at this point, but the situation raises the possibility that there may be other reasons why Pent participates in wing disc development. To this end, it may be important to remember the additional phenotype displayed by *pent* mutants in the wing: loss of the fifth longitudinal vein (Vuilleumier et al., 2010). Whereas scaling abnormalities might explain mispositioning of a vein, vein loss suggests that Pent may have a function independent from its effect on Dpp gradient scaling.

## ACKNOWLEDGMENTS

We thank Giorgos Pyrowolakis, Tetsuya Tabata, Hiroshi Nakato, and Avital Rodal for generously sharing strains. Many thanks to Michelle Digman, Thomas Schilling, and Rahul Warrior for valuable comments and suggestions. Michelle Digman also provided assistance on FCS and ccRICS techniques. This work was funded by NIH grant P50-GM076516.

## AUTHOR CONTRIBUTIONS

Conceptualization, Y.Z. and A.D.L.; Methodology, Y.Z., Y.Q., W.C., Q.N., and A.D.L.; Software, Y.Z., Y.Q., W.C., and A.D.L.; Formal Analysis, Y.Z., Y.Q., W.C., and A.D.L.; Investigation, Y.Z.; Resources, Q.N. and A.D.L.; Writing – Original Draft, Y.Z., Y.Q., and A.D.L.; Writing – Review & Editing, Y.Z., Y.Q., W.C., Q.N., and A.D.L.; Supervision, Q.N. and A.D.L.; Funding Acquisition, Q.N. and A.D.L.

## DECLARATION OF INTERESTS

The authors declare no competing interests.

## METHODS

### Fly strains

The following flies were used in this study: *brk-Gal4, pent*^*2*^, *pent*^*A17*^, *UAS-Pent, UAS-GFP-Pent, pent*^*A17*^,*tkv*^*strII*^ (generous gifts from Giorgos Pyrowolakis), *tkv*^*a12*^, *tkv*^*Df*^, *ubi-Tkv-HA* (generous gifts from Tetsuya Tabata), *dally*^*80*^, *UAS-Dally, y,w,hsFLP; Act*>*y*>*Gal4,UAS-GFP*.*nls* (X; III) (generous gifts from Hiroshi Nakato), *UAS-Dpp-Dendra2* (Zhou et al., 2012), *UAS-tkvmCherry* (a generous gift from Avital Rodal), *ds-Gal4* (provided by Marcos Nahmad). All other stocks were from Bloomington Drosophila Stock Center.

Genotypes in each figure were: **Figure 1**: *+/UAS- GFP; Act5C-Gal4/+* (A, E); *UAS-GFP/UAS-pentRNAi; Act5C-Gal4/+* (B, E); *UAS-GFP/+; hh-Gal4, tubP- Gal80*^*ts*^*/+* (C, E); *UAS-GFP/UAS-pentRNAi; hh-Gal4, tubP-Gal80*^*ts*^*/+* (D, E); *UAS-GFP/+; Act5C-Gal4/UAS- pentRNAi* (E); *UAS-GFP/UAS-pentRNAi; Act5C- Gal4/UAS-pentRNAi* (E); *UAS-GFP/+; hh-Gal4, tubP- Gal80*^*ts*^*/UAS-pentRNAi* (E); *UAS-GFP/UAS-pentRNAi; hh-Gal4, tubP-Gal80*^*ts*^*/UAS-pentRNAi* (E); wild-type (E). **Figure 2**: *ap-Gal4/pent*^*2*^; *UAS-RFP/UAS-GFP-Pent* (A-A’’, D); *cut-Gal4/pent*^*2*^; *UAS-RFP/UAS-GFP-Pent* (B-B’’, D); *pent*^*2*^*/pent*^*2*^; *hh-Gal4, tubP-Gal80*^*ts*^*/UAS-GFP-Pent* (C-D). **Figure 3**: *hsFLP/+; +/UAS-Pent; Act*>*y*>*Gal4, UAS-GFP/UAS-Pent* (A-I’). **Figure 4:** *brk-Gal4/+; +/UAS-GFP* (A, A’, F); *brk-Gal4/+; +/UAS-GFP; +/UAS- sflRNAi* (B, B’, F); *brk-Gal4/+; +/UAS-GFP; +/UAS-Pent* (C, C’, F); *+/UAS-GFP; Act5C-Gal4/UAS-sflRNAi* (D, D’, F); *brk-Gal4/+; +/UAS-GFP; +/UAS-tkvRNAi* (E, E’, F). **Figure 5:** wild-type (A, I), *+/pent*^*2*^ (B, I), *pent*^*2*^*/ pent*^*2*^ (C,I), *tkv*^*Df*^*/tkv*^*strII*^;*ubi-tkv-HA/+* (D,I), *Act5C-Gal4,dally*^*80*^*/UAS-Dally,dally*^*80*^ (E,I), *tkv*^*strII*^*/tkv*^*a12*^,*ubi-tkv-HA;Act5C- Gal4,dally*^*80*^*/UAS-Dally,dally*^*80*^(F,I), *pent*^*2*^,*tkv*^*Df*^*/pent*^*A17*^,*tkv*^*strII*^; *+/ubi-tkv-HA* (G, I), *pent*^*A17*^,*tkv*^*strII*^*/pent*^*2*^,*tkv*^*a12*^,*ubi-tkv-HA; Act5C- Gal4,dally*^*80*^*/UAS-Dally,dally*^*80*^ (H, I). **Figure 7:** *Act5C- Gal4/UAS-pentRNAi* (D, J); *+/tubP-Gal80ts; Act5C- Gal4/UAS-pentRNAi* (A-C, E-H, J); wild-type (I, J).

### Clonal analysis

The *Act*>*y*>*Gal4* transgene was used to generate random GFP marked “flip-out” clones, which were induced by heat-shock of second instar larvae (48-72 hours after egg laying) at 37°C for 20 min, and larvae were allowed to grow at 25°C.

### Egg-laying assay

Flies were kept in a 25°C incubator and eggs were collected on apple juice agar plates with yeast paste at the center. Prior to egg collection, we treated flies with CO_2_ and then let them lay eggs on a plate for 1 h to get rid of old eggs. After that, the flies were transferred to a new plate and the eggs were collected for 1 h. Then collection plates were kept in a 25°C incubator and larvae from these plates were dissected at different hours after egg-laying (AEL).

### Conditional knockdown of *pent*

Eggs from appropriate crosses were collected as above. These eggs were incubated at 30°C (*pentRNAi* on) for 24 h, 48 h or 72 h after egg laying, before being switched to 18°C, which would turn *pentRNAi* off at approximately early first instar stage, early second instar stage or early third instar stage, respectively. The larvae from these eggs then were dissected at the wandering stage. The larvae from the same crosses were raised at 18°C (*pentRNAi* off) and then switched to 30°C (*pentRNAi* on) for 24 h, 48 h, 72 h or 96 h; at that time only wandering larvae were selected for dissection. This results in *pentRNAi* having been on approximately from early-mid third instar stage, late second-early third instar stage, late first-early second instar stage or first instar stage, respectively.

### Antibodies and immunostaining

For immunostaining, larvae were dissected in ice cold phosphate-buffered saline (PBS) and transferred directly into fix solution (4% paraformaldehyde and 0.05M EGTA in PBS). Samples were fixed for 30 min at room temperature. Samples were then washed extensively 5 times for 10 min each with PBT (0.1% Tween-20 in PBS) and blocked overnight at 4°C in blocking solution (1% BSA, 0.3% Deoxycholate and 0.3% TritonX-100 in PBS). Afterwards, samples were incubated with primary antibodies diluted in blocking solution at 4°C overnight then washed 6 times in PBT for 10 min each, and incubated with secondary antibodies and DAPI diluted in PBT for 1.5 h at room temperature on a rotor. After 5 washes in PBT for 10 min each, stained discs were mounted on slides.

The following primary antibodies were used: rabbit anti-pSmad3 (Abcam, EP823Y) 1:1000; mouse anti-Ptc (DSHB, Apa1) 1:1000; mouse anti-Wg (DSHB, 4D4) 1:1000; mouse anti-Dlp (DSHB, 13G8) 1:1000; mouse anti-β-galactosidase (Promega, Z378B) 1:1000. Alexa Fluor-conjugated anit-rabbit and anti-mouse secondary antibodies were used at 1:1000. 100 μg/mL DAPI were diluted 1:1000 in PBT.

### Imaging and image analysis

Wing discs were dissected from larvae of various stages and mounted in ice cold PBS on slides. The slides were kept on ice for live imaging of *Drosophila* wing discs. Images of both fixed and live wing discs were obtained with a Zeiss LSM 780 laser scanning confocal fluorescence microscope.

For imaging of *Drosophila* adult wings, flies were preserved in 70% ethanol. After the 70% ethanol was removed, the wings were plucked and mounted in 50% Canada balsam in xylene on slides. Images of wings were obtained with a Zeiss SteREO Discovery.V8 stereomicroscope.

Images were analyzed using Fiji software. They were quantified by scanning along a straight line starting at the anterior/posterior (A/P) boundary and ending at the edge of the wing pouch, parallel to the anterior-posterior axis, with about 20% dorsal offset from dorsal/ventral (D/V) boundary. Both pMad and Ptc fluorescence intensity profiles were extracted along the line. Automated algorithms were developed in Wolfram *Mathematica* to detect the location of the A/P compartment boundary (the edge of Dpp production region) using these fluorescence intensity profiles. To make sure that the locations of the A/P compartment boundaries determined by the automated algorithms were accurate, we manually checked the intensity profiles of each wing disc, and made corrections if necessary. Posterior compartment size was quantified by following a segmented line along the Wg staining at the D/V boundary (starting from the A/P boundary and ending at the edge of the wing pouch), and measuring the length of the line. To obtain a value of *λ*_app_, the pMad profile was fitted to the function *y*(*x*) = *a e*^-*x*/*λ*^ + *b*, where *λ* = *λ*_app_ and *b* is background intensity, using the “NonlinearModelFit” (non-linear fitting function) in Wolfram *Mathematica*. Similar methods were used to analyze GFP-Pent and DppDendra2 profiles.

### Fitting scaling dynamics

Inspection of pMad gradients at multiple stages of development suggested that *λ*_app_ increases more or less linearly with disc size until a threshold size is reached; after that, *λ*_app_ either remains constant or increases linearly but at a slower rate. To quantify this behavior with a minimum of parameters, we identified a general functional form that transitions sharply from one constant slope to another at a particular point. Briefly, the function is the solution to the differential equation *f′*(*x*) = (*b*- *a*)/(1+(*x*/*c*)^*n*^)+*a*, where *c* represents the switching point, *b* is the slope when *x*<<*c* and *a* is the slope when *x*>>c (the solution can be represented explicitly as a hypergeometric function). The Hill coefficient, *n*, controls the sharpness of the switching at the turning point *c*, and for sufficiently large values of *n* has almost no influence on the quality of the fit of this function to the data. In the plots in Figures 4-5, fitting was carried out using “NonlinearModelFit” in Wolfram *Mathematica*, and a value of *n*=9.

### Mathematical modeling

Mathematical modeling was performed using MATLAB and *Mathematica*. For details, see Supplemental Information.

### Fluorescence correlation spectroscopy (FCS) and cross-correlation raster image correlation spectroscopy (ccRICS)

Single point FCS assay (Zhou et al., 2012) and ccRICS assay (Digman et al., 2013) were performed using an Olympus FluoView FV1000 confocal microscope with a 60x/1.2 water immersion objective. Data were analyzed with SimFCS software (Laboratory for Fluorescence Dynamics, University of California, Irvine). Each FCS measurement lasted for 100 seconds. To collect ccRICS data, the following settings were applied: pixel size 0.077 μm, pixel dwell time 10 μs, line time 3.83 ms.

---

**Figure S1:**
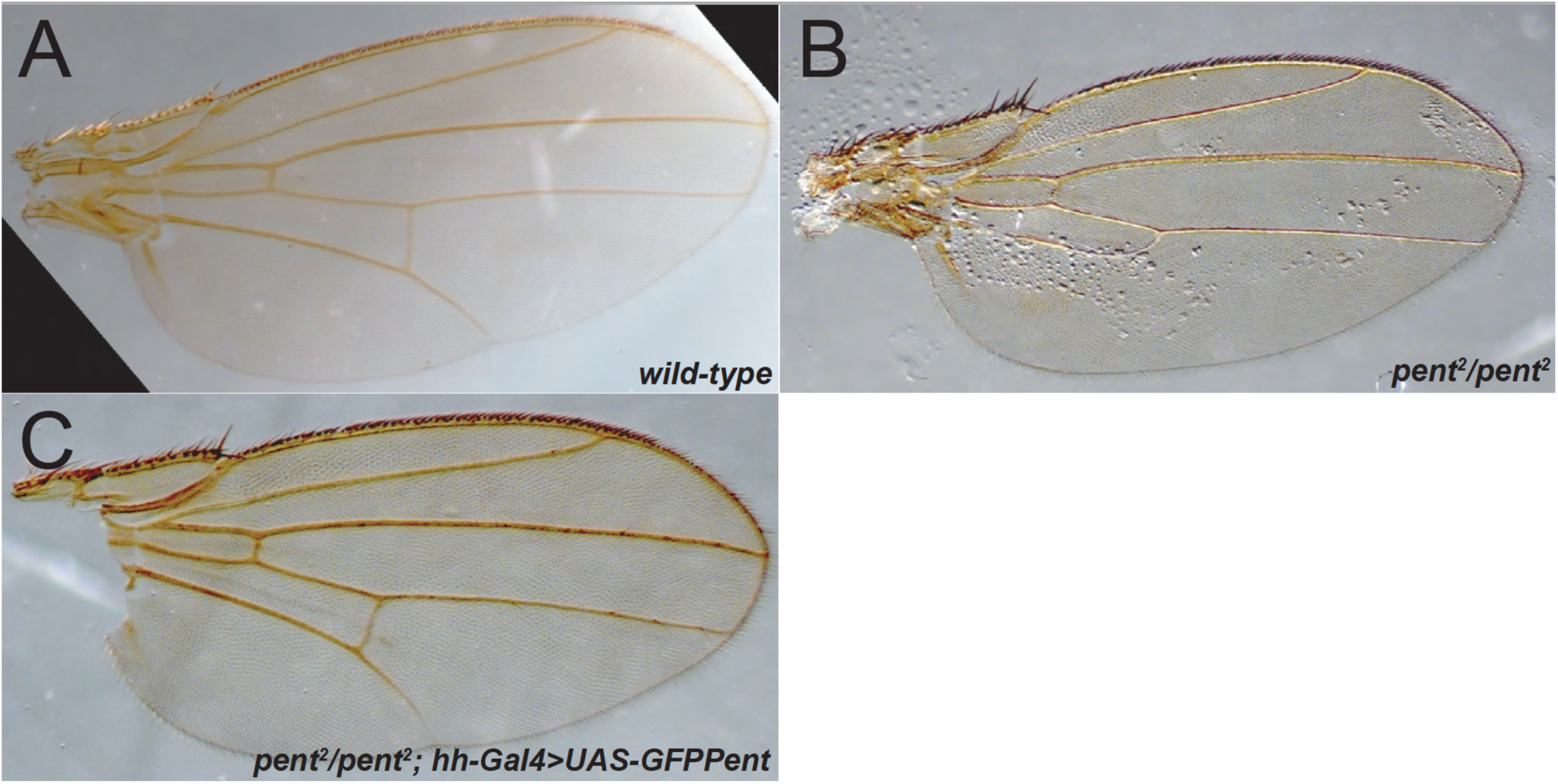
GFP-Pent is functional. (A-C) Comparison of wild-type (A), *pent*^*2*^*/pent*^*2*^ (B) and *pent*^*2*^*/pent*^*2*^; *hh-Gal4/UAS-GFP-Pent* (C) wings. The *Pent* mutant phenotype in the adult posterior wing (loss of longitudinal vein 5) is rescued by overexpression of *GFP-Pent* in the posterior compartment of the wing disc.

**Figure S2:**
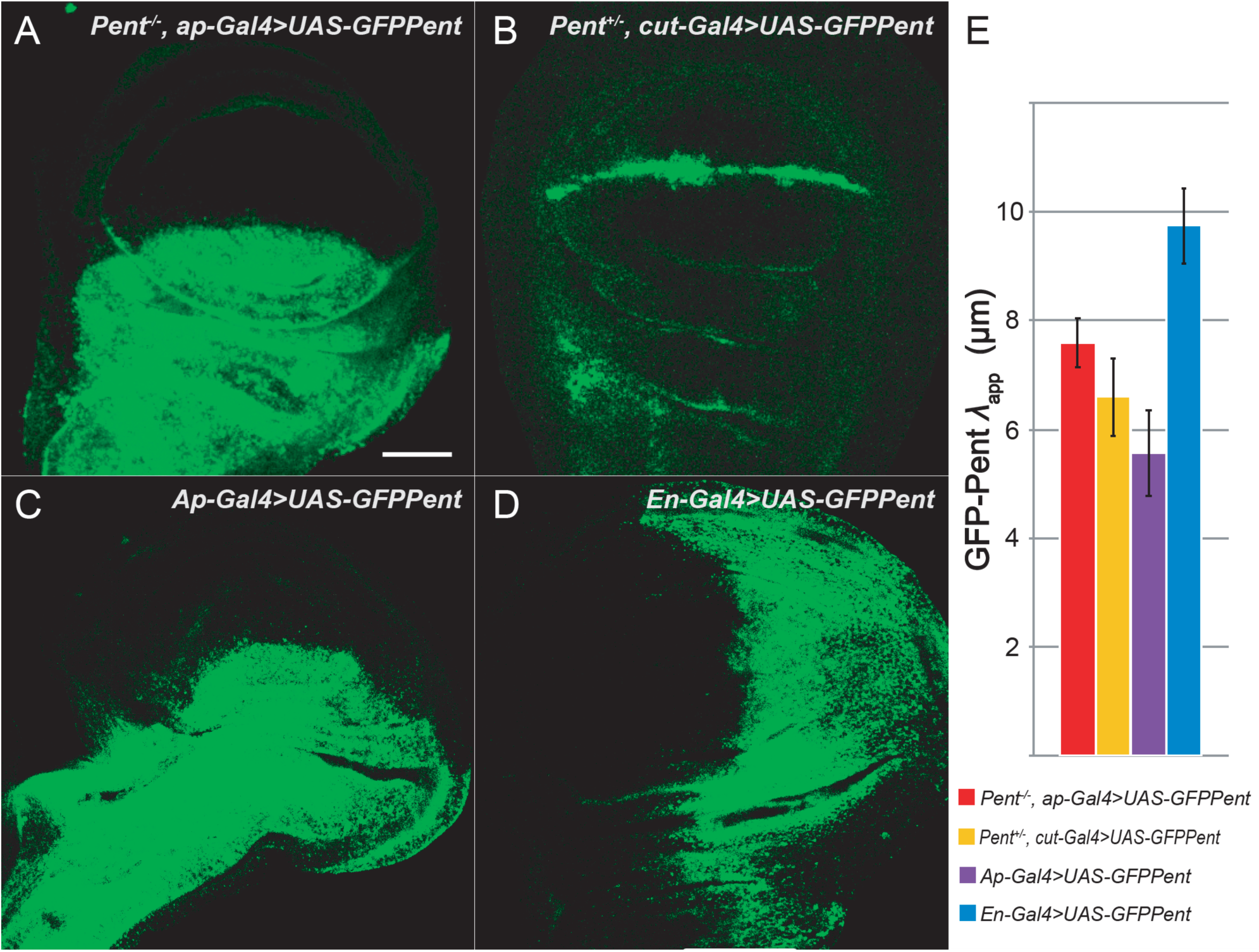
Quantification of GFP-Pent spreading under additional conditions (A-D) Representative wing discs of *pent*^*2*^*/pent*^*2*^, *ap-Gal4; UAS-GFP-Pent/+* (A), *pent*^*2*^*/cut-Gal4; UAS-GFP-Pent/+* (B), *+/ap-Gal4; UAS-GFP-Pent/+* (C), and *+/en-Gal4; UAS-GFP-Pent/+* (D). (E) *λ*_*app*_ for GFP-Pent in these four genotypes is similar (<10 µm) to that shown in Fig. 2. The scale bar is 50 µm.

**Figure S3:**
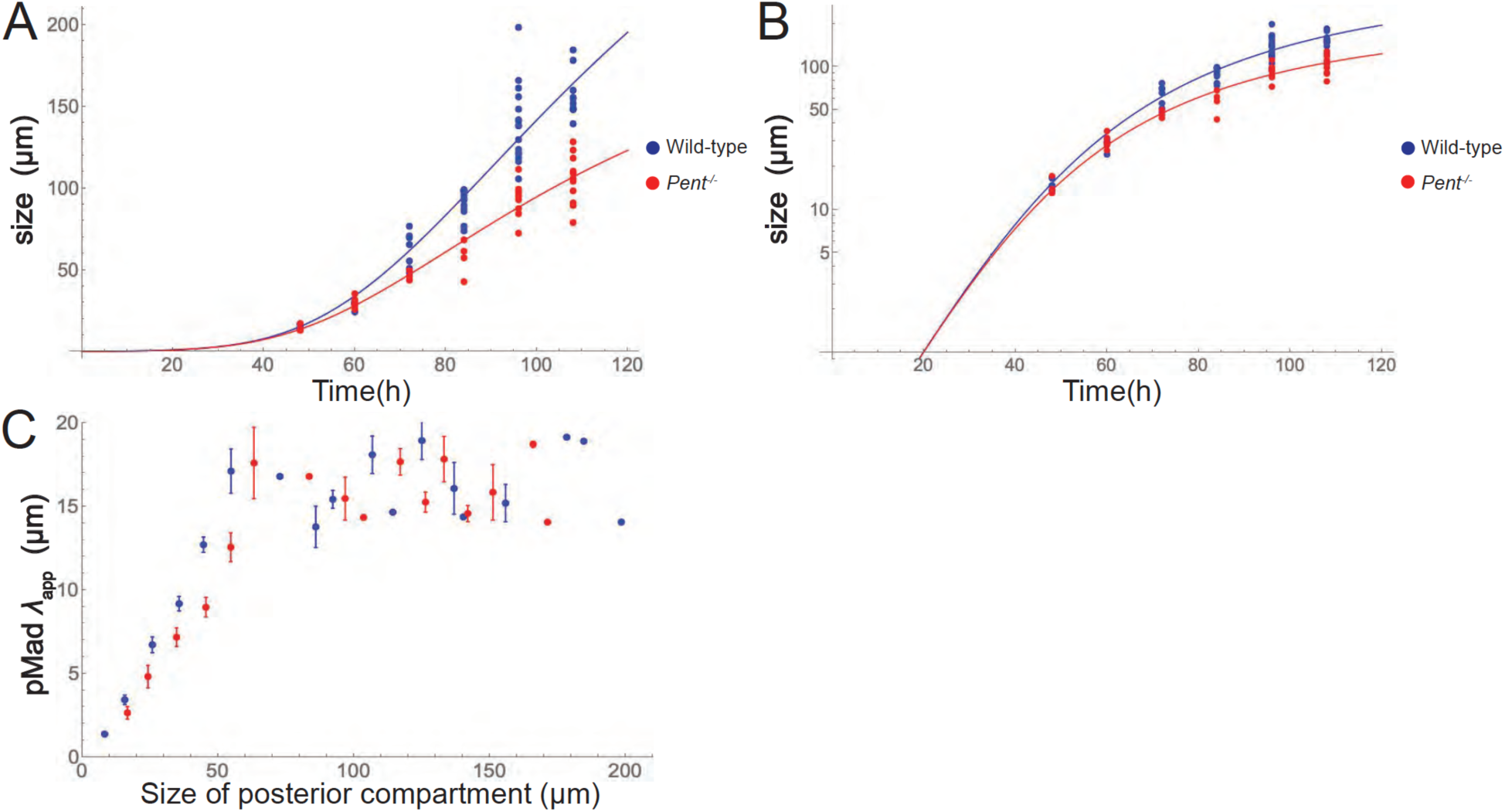
Dynamics of growth and pMad *λ*_*app*_ as a function of time, genotype and measurement algorithm. (**A-B**) The size of the posterior compartment was measured in wild-type (blue) and *pent* mutant (red) wing discs at various times after egg laying (72 h, 84 h, 96 h, 108 h, 120 h and 132 h AEL) and plotted on both linear and logarithmic scales. Note that growth starts off close to exponential but gradually slows. In addition, *pent* mutant discs grow more slowly than wildtype. Data are fit to a smooth curve as described in Supplemental Material. The coefficients of determination (R^2^) for wildtype and *pent*^*-/-*^ are 90.6% and 90.3%, respectively. (**C**) Morphogen gradient scaling is a discontinuous process. pMad *λ*_*app*_ was measured in the posterior compartments of 77 wildtype discs at a variety of stages. To facilitate direct comparison with previous studies, we quantified compartment size in two different ways, either measuring the length of a segmented line tracing Wg antibody staining at the D/V compartment boundary, starting at the A/P compartment boundary and ending at the edge of the wing pouch edge (blue symbols), or taking the length of a straight line along the D/V compartment boundary, starting from the A/P compartment boundary and ending at the edge of the wing disc (blue symbols). The first compares with that used by Hamaratoglu et al. (Hamaratoglu et al., 2011), while the second is the one that was used by Wartlick et al. (Wartlick et al., 2011). In both cases, the data were binned by compartment size into 10 µm bins. Averages ± SEM are plotted for each bin. The results agree with both studies, except that, by analyzing a larger number of large discs, we can observe a sharp discontinuity in scaling after posterior compartments reach about 60 µm in size.

**Figure S4:**
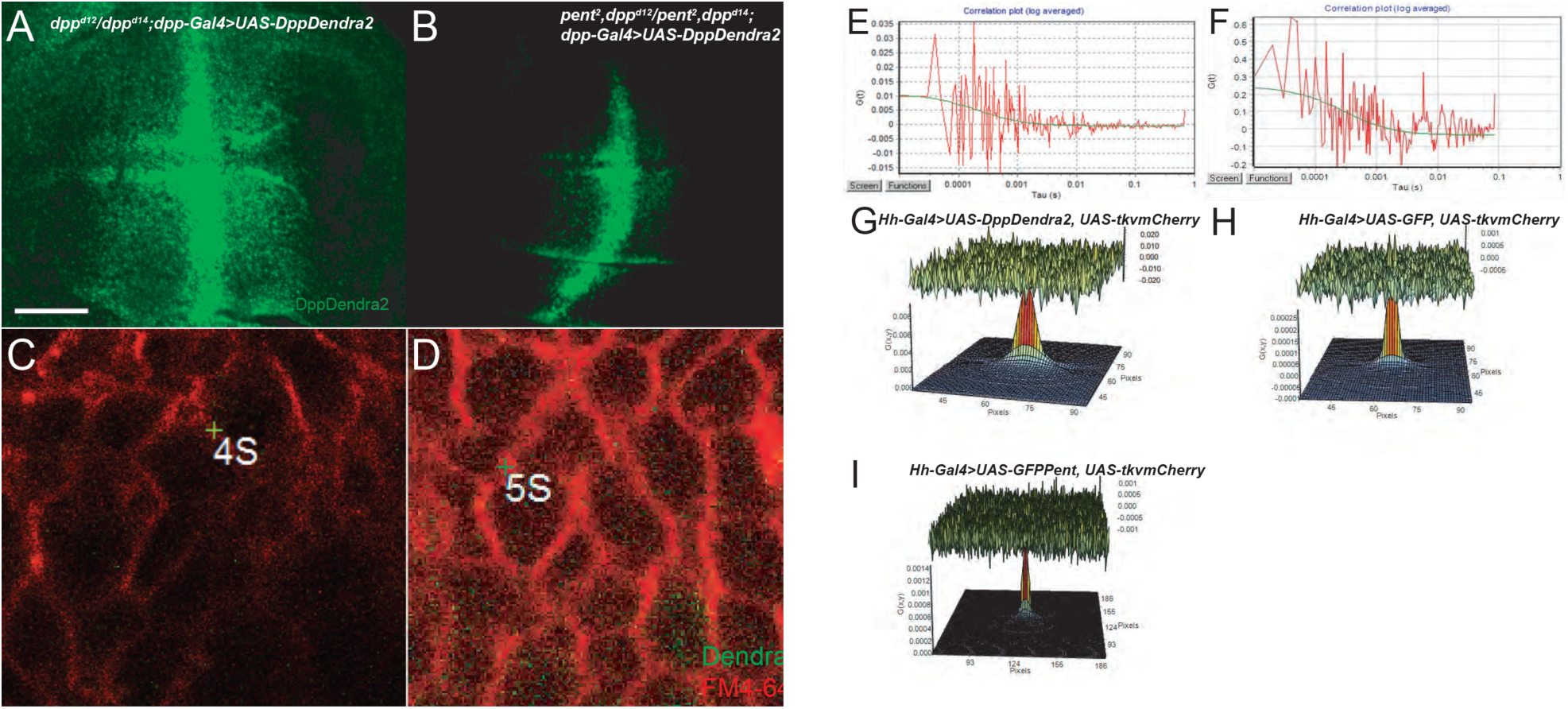
Dynamics of Dpp diffusion and gradient formation in wing discs of *pent* mutants. (**A-B**) To visualize Dpp, endogenous Dpp was replaced with DppDendra2 (Zhou et al., 2012). Representative images of wing discs of *dpp*^*d12*^*/dpp*^*d14*^; *dpp-Gal4/UAS-DppDendra2* (A) and *pent*^*2*^, *dpp*^*d12*^*/ pent*^*2*^, *dpp*^*d14*^; *dpp-Gal4/UAS-DppDendra2* (B) (Green: DppDendra2). The average decay length of DppDendra2 was found to be 21±1.81 µm in wild-type discs (n=8) and 12±0.96 µm in *pent* mutants (n=8). Scale bar = 50 µm. (**C-F**) Pent does not affect Dpp diffusivity. High magnification images of wing discs of genotypes in panels A and B are shown in C and D, respectively. Plasma membranes were visualized with FM4-64 staining (red) and Dendra2 fluorescence in green. Intercellular locations where single-point fluorescence correlation spectroscopy (FCS) was performed are marked by reticles [the labels “4S” and “5S”, which are placed on the image by the microscopy software, are unimportant]. Fluorescence autocorrelation functions for the genotypes in panels C and D are shown in panels E-F, respectively. By obtaining and fitting such data for multiple samples, we obtain an average Dpp diffusion coefficient of 21±0.66 µm^2^/s for the control group (*n*=4) and 21±1.01 µm^2^/s for *pent* mutants (n=6). These values are in close agreement with those measured previously for wildtype *DppDendra2-*expressing discs (Zhou et al., 2012). (**G-I**) Testing for direct interactions between Pent and Tkv. Cross-correlation raster-scanning intensity correlation spectroscopy is a method that uses fluorescence co-fluctuation to infer co-diffusion of molecular species, and hence binding between species. Wing discs were engineered to express mCherry-labeled Tkv in the posterior compartment (Deshpande et al., 2016), together with either *DppDendra2* (G), GFP (H) or GFP-Pent (I). As each pixel is illuminated, fluorescence co-fluctuation is assessed at every other pixel. Data are aligned so that the illuminated pixel is centered of each frame and then summed. Peak heights report the degree of cross-correlation (note different axis values in G, H and I) and peak widths reflect diffusion of the cross-correlated species. The broad and tall peak (maximum G ∼0.008) in G likely reflects binding of DppDendra2 to Tkv. The narrow and low peak in I (maximum G ∼0.0014) is consistent with an absence of binding (as expected for GFP and Tkv). The peak in H is similar to that in I, suggesting that there is no significant interaction between Pent and Tkv.

**Figure S5:**
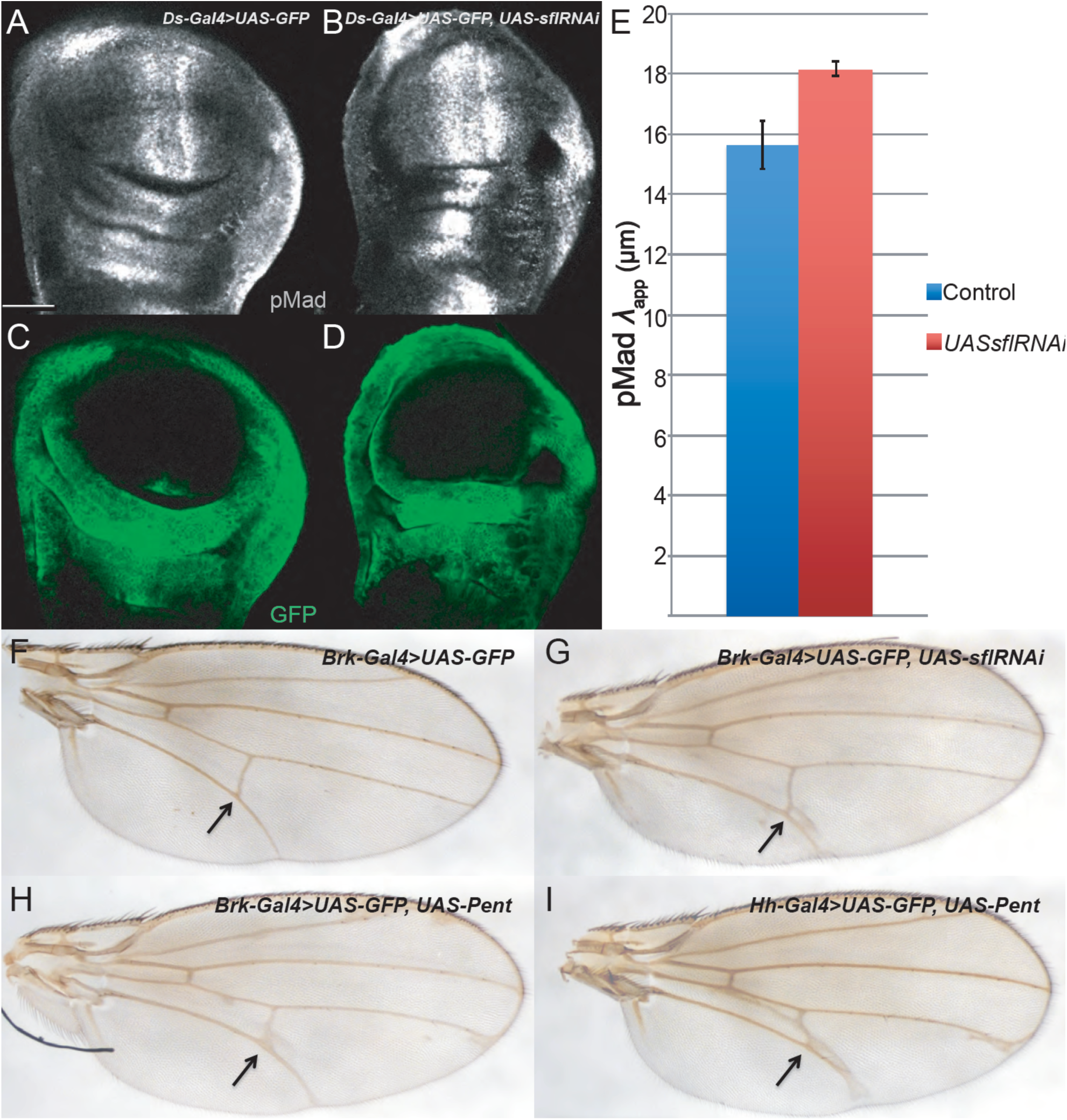
Locally disabling Dpp coreceptors phenocopies *pent* overexpression. (**A-D**) Inhibiting *sfl* function in the Dachsous (ds) expression domain—which is similar to the Pent expression domain—enhances Dpp spreading. Representative wing discs of *ds-Gal4/UAS-GFP* (A, C) and *ds-Gal4/UAS-GFP; +/UAS-sflRNAi* (B, D) are shown; GFP fluorescence is visualized in green pMad staining in. (**E**) Quantification of pMad *λ*_*app*_ in the corresponding flies shows that *λ*_*app*_ increases when coreceptor are disabled in the *ds*-domain. Scale bar = 50 µm. (**F-I**) Adult wing phenotypes. Genotypes are *brk-Gal4; UAS-GFP* (F), *brk-Gal4; UAS-GFP; UAS-sflRNAi* (G), *brk-Gal4; UAS-GFP; UAS-pent* (H), and *UAS-GFP; hh-Gal4/UAS-pent* (I). Posterior cross veins are pointed out by arrows. *Pent*-overexpression produces marked thickening in the vicinity of the posterior crossvein, which is phenocopied by expressing *sflRNAi* in the *brk* domain.

**Figure S6:**
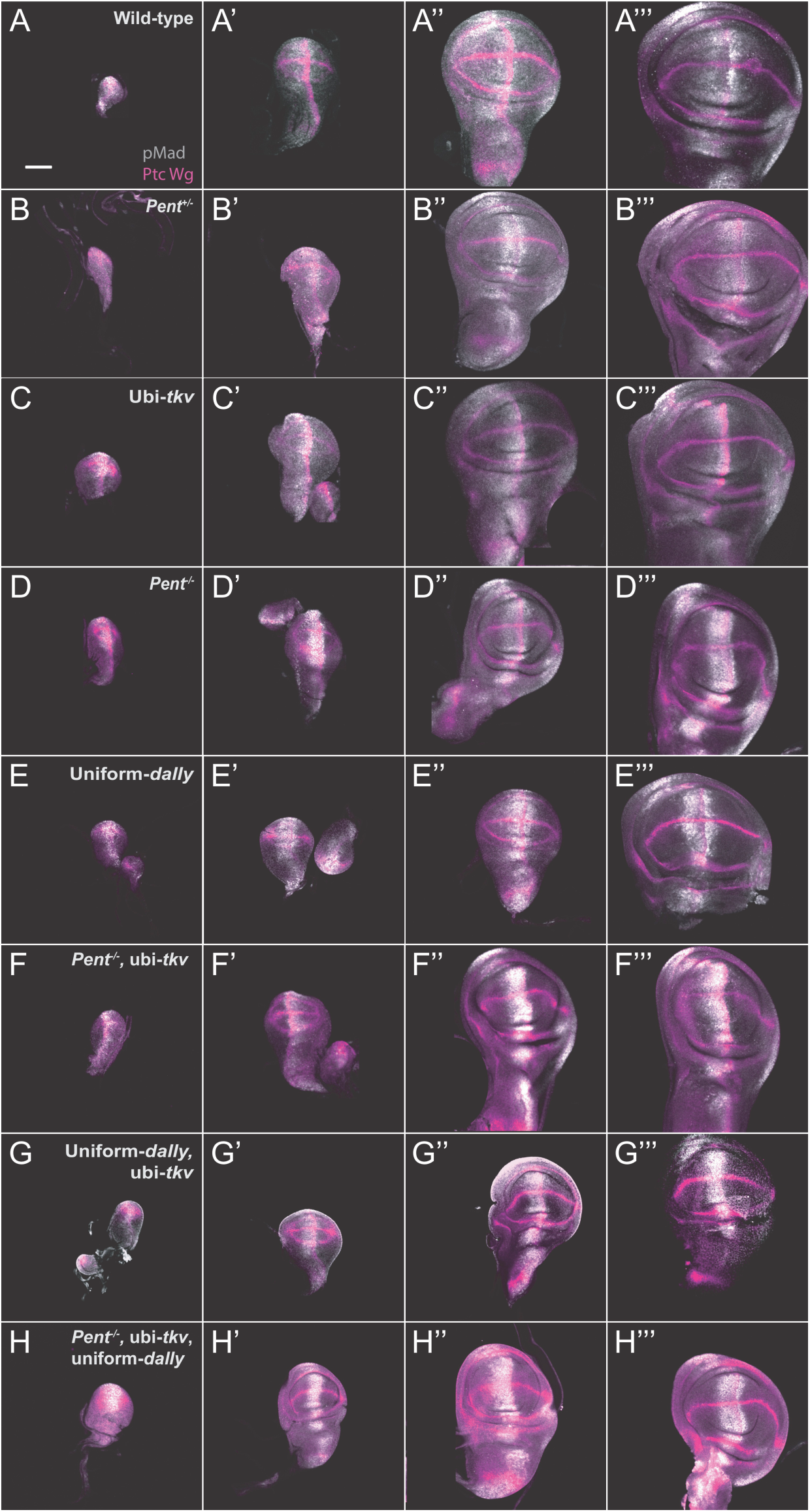
Representative images of wing discs of wild-type (A-A’’’), *+/pent*^*2*^ (B-B’’’), *tkv*^*Df*^*/tkv*^*strII*^; *ubi-tkv-HA* (C-C’’’), *pent*^*2*^*/pent*^*2*^ (D-D’’’), *Act5C-Gal4, dally*^*80*^*/UAS-Dally, dally*^*80*^ (E-E’’’), *pent*^*2*^, *tkv*^*Df*^*/pent*^*A17*^, *tkv*^*strII*^; *ubi-tkv-HA* (F-F’’’), *tkv*^*strII*^*/tkv*^*a12*^, *ubi-tkv-HA; Act5C-Gal4, dally*^*80*^*/UAS-Dally, dally*^*80*^ (G-G’’’) and *pent*^*A17*^, *tkv*^*strII*^*/pent*^*2*^, *tkv*^*a12*^, *ubi-tkv-HA; Act5C-Gal4, dally*^*80*^*/UAS-Dally, dally*^*80*^ (H-H’’’) in different stages stained with anti-pMad, anti-Ptc and anti-Wg antibodies (gray, magenta). Scale bar = 50 µm.

**Figure S7:**
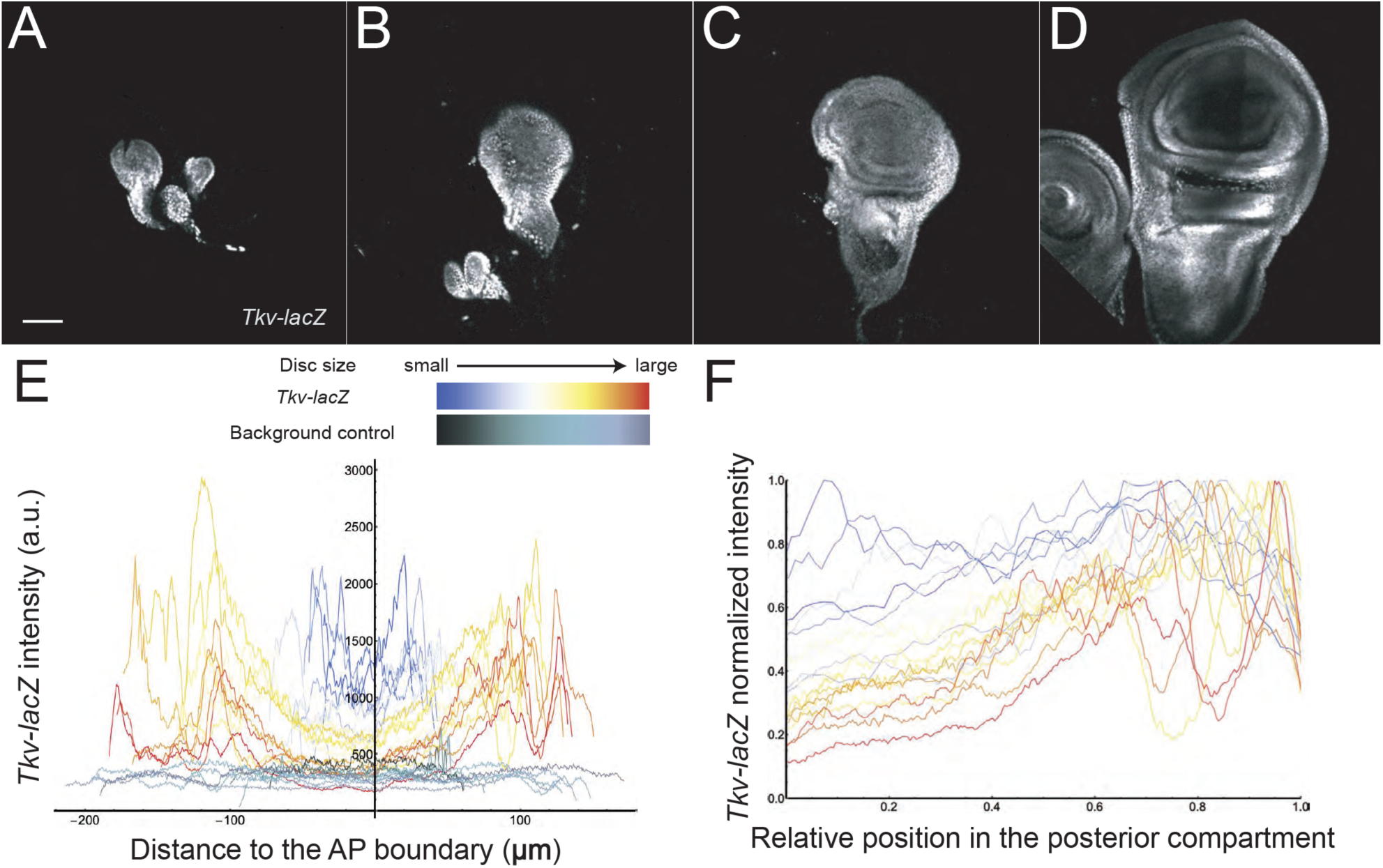
Repression of Tkv in the center of the wing pouch builds gradually over the course of wing disc development. *Tkv* expression was monitored using the enhancer trap line, *tkv-lacZ*. (**A-D**) Representative images of wing imaginal discs of *tkv-lacZ* at different developmental stages, stained with anti-β-galactosidase antibodies. Scale bar = 50 µm. **(E)** *Tkv* expression profiles were extracted along the D/V boundary for discs of multiple sizes. Plots are color-coded by disc size (blue for early, small discs to red for late, large discs. (**F**) Profiles in panel E were normalized to maximal *tkv* expression level and posterior compartment size. The data indicate that the degree of tkv repression grows substantially as discs get larger, consistant with amplitude growth of the Dpp gradient.

## Notes

#### Summary of Updates

We identified a typo in one of the equations in the last section of the Supplemental Information, and have corrected this.

