## Supplemental Information for "Feedback control of morphogen gradient scale"

### Section 1. Description of the mathematical model

The conceptual network and partial differential equations used to model the Dpp gradient on a growing wing disc are shown below. Arrows depict biochemical interactions; lines ending in bars represent regulatory inhibition, and lines ending in circles regulatory activation. The domain modeled is the intercellular (basolateral) spaces of the posterior compartment of the wing disc, which is represented as a one-dimensional reaction-diffusion system. The one-dimensional approximation assumes that morphogen flux in the dorsoventral and apicobasal directions is negligible. This is probably a good assumption at large disc sizes, but may be less so at small size (cf. Lander et al., 2011). We model the basal rates of synthesis of gene products as constant in time and space, as modified by pMad or Brk, except in the following cases: we lower Tkv production and raise coreceptor production in the Dpp-production region, to capture known effects of Hedgehog signaling in that region (Tanimoto et al., 2000). In addition, we model the basal rate of Brk synthesis as continuously increasing during disc growth, in order to fit the data of (Hamaratoglu et al., 2011), who show that peak Brk levels rise more than 10-fold over the course of wing disc development (because peak Brk expression occurs where Dpp signaling is essentially negligible, such changes cannot be attributed to an effect of Dpp). We model growth of wildtype discs to fit our own observations of disc growth rate [Figure S3], which are similar to those published by (Wartlick et al., 2011). For some genotypes, including homozygous *pent* mutants, we adjusted the growth rate so that discs finish growing at a smaller size (see Section

2, below), in accordance with published data on *pent* discs (Ben-Zvi et al., 2011; Vuilleumier et al., 2010), as well as our own observations [Fig. 5 and S6].

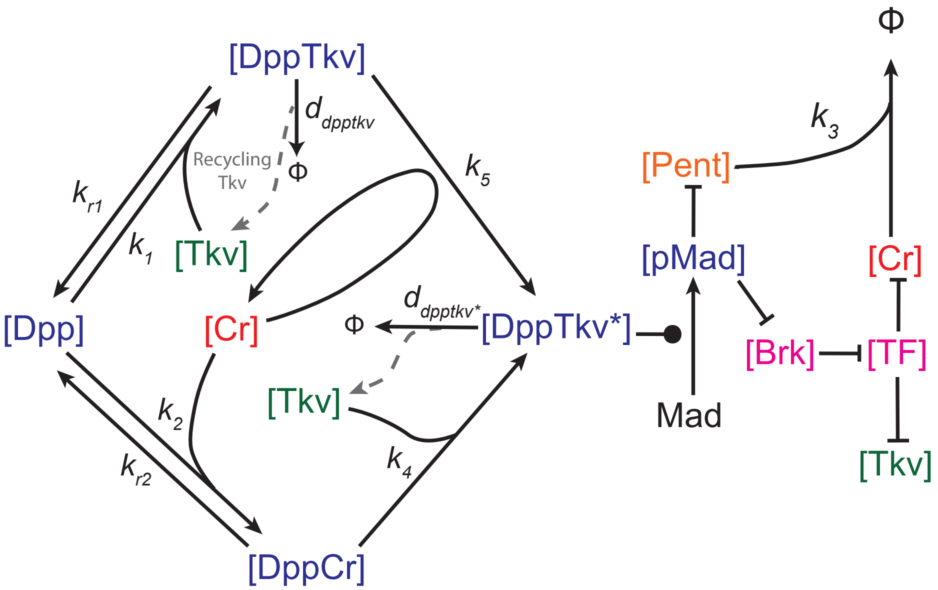


$$
\begin{aligned}
&\frac{\partial[Dpp]}{\partial t}+\frac{\partial\left(V[Dpp]\right)}{\partial x}=\frac{v_{dpp}}{1+\left(\frac{x}{prod(t)}\right)^{20}}+D_{dpp}\Delta[Dpp]-k_1[Dpp][Tkv]+k_{r1}[Dpp-Tkv]-k_2[Dpp][Cr]+k_{r2}[Dpp-Cr]-\frac{d_{dpp}}{1+\left(\frac{x}{prod(t)}\right)^{20}}[Dpp],\\
&\frac{\partial[Tkv]}{\partial t}+\frac{\partial\left(V[Tkv]\right)}{\partial x}=v_{tkv}-k_1[Dpp][Tkv]+k_{r1}[Dpp-Tkv]-k_4[Tkv][Dpp-Cr]+d_{dpptkv}[Dpp-Tkv]+d_{dpptkv^*}[Dpp-Tkv^*]-d_{tkv}[Tkv],\\
&\frac{\partial[Cr]}{\partial t}+\frac{\partial\left(V[Cr]\right)}{\partial x}=v_{cr}-k_2[Dpp][Cr]+k_{r2}[Dpp-Cr]+k_4[Tkv][Dpp-Cr]-k_3[Pent][Cr]-d_{cr}[Cr],\\
&\frac{\partial[Dpp-Tkv]}{\partial t}+\frac{\partial\left(V[Dpp-Tkv]\right)}{\partial x}=k_1[Dpp][Tkv]-k_{r1}[Dpp-Tkv]-k_5[Cr][Dpp-Tkv]-d_{dpptkv}[Dpp-Tkv],\\
&\frac{\partial[Dpp-Tkv^*]}{\partial t}+\frac{\partial\left(V[Dpp-Tkv^*]\right)}{\partial x}=k_5[Cr][Dpp-Tkv]+k_4[Tkv][Dpp-Cr]-d_{dpptkv^*}[Dpp-Tkv^*],\\
&\frac{\partial[Dpp-Cr]}{\partial t}+\frac{\partial\left(V[Dpp-Cr]\right)}{\partial x}=k_2[Dpp][Cr]-k_{r2}[Dpp-Cr]-k_4[Tkv][Dpp-Cr]-d_{dppcr}[Dpp-Cr],\\
&\frac{\partial[pMad]}{\partial t}+\frac{\partial\left(V[pMad]\right)}{\partial x}=v_{pmad}[Dpp-Tkv^*]-d_{pmad}[pMad],\\
&\frac{\partial[brk]}{\partial t}+\frac{\partial\left(V[brk]\right)}{\partial x}=\frac{x_{\max}v_{brk}}{\left(1+\left(\frac{[pMad]}{EC_{brk}}\right)^2\right)}-d_{brk}[brk],\\
&\frac{\partial[TF]}{\partial t}+\frac{\partial\left(V[TF]\right)}{\partial x}=\frac{v_{tf}}{1+\left(\frac{[brk]}{EC_{TF}}\right)^2}-d_{tf}[TF],\\
&\frac{\partial[Pent]}{\partial t}+\frac{\partial\left(V[Pent]\right)}{\partial x}=\frac{v_{pent}}{1+\left(\frac{[pMad]}{EC_{pent}}\right)^2}-k_3[Pent][Cr]-d_{pent}[Pent].\\
&v_{tkv}=tkv1+\frac{tkv2}{1+\left(\frac{[TF]}{EC_{tkv}}\right)^2}+\frac{tkv3}{1+\left(\frac{x}{prod(t)}\right)^{-20}},\quad v_{cr}=cr1+\frac{cr2}{1+\left(\frac{[TF]}{EC_{Cr}}\right)^2}+\frac{cr3}{1+\left(\frac{x}{prod(t)}\right)^{20}}.
\end{aligned}
\tag{1}
$$

In the above system of equations, $[P](x,t)$ denotes the concentration of species $P$ at location $x$ and at time $t$. The spatial domain $[0, x_{max}(t)]$ represents the region of posterior compartment. To represent disc growth, $x_{max}$ increases according to: $x_{max}(t) = x_0 e^{at\,{}_2F_1\left(1,\frac{1}{n},1+\frac{1}{n},-bt^n\right)}$ (derivation in Section 2). Dpp is the only diffusive species in this model with diffusion term $D_{dpp}\Delta[dpp]$ and diffusion coefficient $D_{dpp}$. The term $\frac{\partial(V[P])}{\partial x}$ can be split into two terms $[P]\frac{\partial V}{\partial x}$ and $V\frac{\partial[P]}{\partial x}$ representing dilution and advection driven by disc growth, respectively. $V(x,t)$ is the disc growth velocity at location $x$ and its value at $x_{max}$ represents for the growth rate of the entire posterior compartment: $V(x_{max},t) = \frac{dx_{max}}{dt}$. We assume the disc grows homogeneously over the entire space, and $V(x,t)$ is a linear function of x:

$$V(x,t) = \frac{x}{x_{max}(t)} V(x_{max},t). \qquad (2)$$

$d_P$ is the degradation rate of $P$. $k_i[P][Q]$, $(i=1,2,3,4,5)$, are association rates between P and Q, and $k_{ri}[PQ]$ ($i$=1,2) are dissociation rates of the complex formed by P and Q. We assume that Dpp is synthesized in a localized source (termed the production region), and the size of Dpp production region grows at the same rate as the rest of posterior compartment. Specifically, we take $prod(t) = p * x_{max}(t)$, with $p = 0.12$. Dpp production is then modeled by $v_{dpp}/(1+(x/prod(t)^{20}))$. The high-exponent Hill function essentially approximates a step function. A Dpp degradation term is also added in the production region: $d_{dpp}/(1+(x/prod(t)^{20}))$.

The production rate of Tkv, $v_{tkv}$, contains three terms: $tkv1$ is a base production rate in the entire disc; $tkv2/(1+([TF]/EC_{tkv})^2)$ represents the production regulated by TF, which stands for downstream transcription factors repressed by Brinker (Brk); Tkv synthesis is low inside the Dpp production region due to the effect of Hedgehog (Tanimoto et al., 2000) and $tkv3/(1+(x/prod(t))^{-}$

$^{20}$) is used to model the additional Tkv production outside of the production region. Dally and Dlp are lumped together as "Co-receptor" (Cr) in this model. The production rate of co-receptor, $v_{cr}$, contains three terms: $cr1$ is base production rate; $cr2/(1+([TF]/EC_{Cr})^2)$ represents the production regulated by TF; and because co-receptor synthesis is high inside Dpp production region, due to effect of Hedgehog (Tanimoto et al., 2000), $cr3/(1+(x/prod(t))^{20})$ is used to model the addition Cr production in the Dpp production region.

To model the production of Brinker (Brk), which is repressed by pMad, we multiply a basal production rate by $1/(1+ [pMad]/EC_{brk})^2)$, however, because basal Brk production appears to increase markedly with disc size (Hamaratoglu et al., 2011), we take the basal production rate to be a constant $v_{brk}$ times $x_{\max}$ (disc diameter). To model the production of TF, we multiply a basal production rate $v_{tf}$ by $1/(1+ [Brk]/EC_{TF})^2)$. To model the production of Pent, we multiply a basal production rate $v_{Pent}$ by $1/(1+ [pMad]/EC_{Pent})^2)$.

The biochemical steps in the assembly of the active form of the Dpp receptor are modeled to reflect that fact that TGF-beta family receptors assemble in a two-stage process which, for the BMP branch of the TGF-beta family usually involves initial binding to type I receptors (e.g. Tkv) and subsequent recruitment of type II receptors. Thus, the species DppTkv may be construed to represent complexes that lack type II receptors while the species DppTkv* represents complexes containing both type I and II receptors.

We model co-receptor activity according to the results of (Kuo et al., 2010), who showed that HSPGs catalyze the conversion of BMP-type I receptor complexes into BMP-type I receptor-type II receptor complexes. Rate constant $k_5$ captures this behavior. At the same time, because Dpp can bind HSPGs, we also model direct reversible binding, and allow for the possibility that Dpp initially bound to HSPGs can also recruit type I and type II receptors; this latter behavior is captured by $k_4$, but as described later, the value of $k_4$ may be set effectively to zero without having significant effect on the model output.

In solving system (1) over time and space it is necessary to specify initial conditions for all variables and boundary conditions for the one diffusing species Dpp. The boundary conditions are no-flux at x=0, i.e. $\left.\frac{d[Dpp]}{dx}\right|_{x=0} = 0$, and absorbing at $x_{\max}$, i.e. $[Dpp]\big|_{x=x_{\max}(t)} = 0$. The no-flux condition is justified by the symmetry of the problem (Dpp feeds gradients in both anterior and posterior compartments), and the absorbing condition creates a sink a $x$=$x_{\max}$.

The initial posterior compartment size is taken to be 0.1 μm—smaller than the actual size of discs—in order to provide sufficient simulation time for results to become independent of initial conditions. The initial conditions are then obtained by running the simulation in the fixed initial domain for 10 hours starting from zero values for all species. We verified that these conditions produced results that were independent of initial condition choices.

The total simulation time is 120 hours. We take time=0 to correspond to 24 hours after egg laying, which is consistent with the convention adopted by (Wartlick et al., 2011).

**Section 2. Derivation of the Growth Rate Function**

The rate at which discs grow is not constant, but slows as larval development proceeds. To determine what growth rate function to use in the model, we measured compartment sizes experimentally (Fig. S3). To fit those data to a simple equation we considered the following function which describes an arbitrary system that is growing exponentially but slowing according to a declining Hill function of time.

$$\begin{cases} \dfrac{dx_{\max}}{dt} = \dfrac{ax_{\max}}{1+bt^n} = f(x_{\max}, t) \\ x_{\max}(0) = x_0. \end{cases} \qquad (3)$$

The general solution to (3) is $x_{\max}(t) = x_0 e^{at\,{}_2F_1\left(1,\frac{1}{n},1+\frac{1}{n},-bt^n\right)}$, where ${}_2F_1$ is the hypergeometric function:

$$ {}_2F_1(a,b;c;z)=\sum_{n=0}^{\infty}\frac{(a)_n(b)_n}{(c)_n}\frac{z^n}{n!}. \tag{4} $$

Here $(q)_n$ is the Pochhammer symbol, which is defined by:

$$ (q)_n=\begin{cases}1, & n=0;\\ q(q+1)\cdots(q+n-1), & n>0.\end{cases} \tag{5} $$

We used the built-in function *NonlinearModelFit* in Mathematica to fit the experimental data (Fig. S3) to the above function. By testing various integers *n*, the best fit was found to be given by *n*=3. We then used this function to describe the growth of $x_{\max}$ over time in the model. Although the mathematical form is different from that proposed by (Wartlick et al., 2011) for the wing disc, the two functions are very similar in shape.

**Section 3. Lagrangian framework for solving mathematical equations**

The spatial domain of Eq. (1) is time-dependent, whereas PDEs solvers usually require a fixed domain. We therefore use the following linear coordinate transformation to transfer the dynamical spatial domain onto a fixed domain:

$$ \begin{cases}x=r(\tau)X\\ t=\tau\end{cases}. \tag{6} $$

where $r(\tau)=\frac{x_{\max}(\tau)}{x_0}$. The transferred spatial domain is $X\in[0,x_0]$, where $x_0$ is the initial posterior compartment size shown in Eq. (3). Derivatives in the Lagrangian coordinate system $(X,\tau)$ have the following relationships to derivatives in the original coordinate system $(x,t)$:

$$ \begin{cases}\frac{\partial}{\partial X}=r\frac{\partial}{\partial x}\\ \frac{\partial^2}{\partial X^2}=r^2\frac{\partial^2}{\partial x^2}\\ \frac{\partial}{\partial \tau}=\frac{\partial}{\partial t}+\frac{\partial}{\partial x}\frac{\partial x}{\partial \tau}=\frac{\partial}{\partial t}+\frac{1}{r}\frac{dr}{d\tau}\frac{\partial}{\partial X}\end{cases} \tag{7} $$

The transformed growth velocity $\tilde{V}(X,\tau)$ has a similar relationship to Eq. (2)

$$\tilde{V}(X,\tau) = \frac{X}{x_0}\tilde{V}(x_0,\tau). \tag{8}$$

Using Eq. (7) and Eq. (8), we have

$$\frac{\partial V}{\partial x} = \frac{1}{r}\frac{\partial \tilde{V}}{\partial X} = \frac{1}{r}\frac{1}{x_0}\tilde{V}(x_0,\tau) = \frac{f(x_{\max}(\tau),\tau)}{x_{\max}(\tau)}. \tag{9}$$

For any equation in Eq. (1) with the general form

$$\frac{\partial [P]}{\partial t} + \frac{\partial \left(V[P]\right)}{\partial x} = F\left([P],x,t\right) + D\Delta[P]. \tag{10}$$

the transformed equation in Lagrangian coordinate is given by

$$\frac{\partial [P]}{\partial \tau} = D\left(\frac{x_0}{x_{\max}}\right)^2 \frac{\partial^2 [P]}{\partial X^2} + F([P],r(\tau)X,\tau) - \frac{f(x_{\max}(\tau),\tau)}{x_{\max}(\tau)}[P]. \tag{11}$$

Since both boundary conditions are homogeneous, the transformed equation inherits the boundary conditions from the original condition: absorbing boundary at one side and no-flux boundary at the other. We solve the transformed equations using PDEs solver *pdepe* in MATLAB 2015b.

**Section 4. Parameter selection**

We modeled eight genotypes in this work: wildtype; $pent^{+/-}$; $pent^{-/-}$; ubi-*tkv*; uniform *dally*; ubi-*tkv*, uniform *dally*; ubi-*tkv*, $pent^{-/-}$; ubi-*tkv*, uniform *dally*, $pent^{-/-}$. Initially, we searched parameter space for sets of parameters that would provide a relatively good fit for the wildtype and $pent^{-/-}$ phenotypes (in the latter case we set Pent production rate and the binding rate ($v_{\text{pent}}$) between Pent and Cr ($k_3$) to zero). Latin Hypercube sampling (Tang, 1993) was used as a random number generator for logarithmic sampling (e.g. $10^{a+b\xi}$ where $\xi$ is the random number) to allow sampling to be uniform across orders of magnitude. Sampled parameter ranges were selected based, where possible, on published results, but more commonly to cover ranges that were biophysically plausible and associated with time scales not substantially longer than the simulation time (in practice this meant sampling ranges in some cases as large as five orders of magnitude). Overall 150,000 independent parameter sets were explored.

The behaviors that the model needed to reproduce were not just scaling of $\lambda_{app}$ for pMad, but also scaling of $\lambda_{app}$ for total Dpp; Dpp and pMad gradient shapes that are close to exponential at the end of larval development; degrees of central-suppression of Tkv and Cr at the end of larval development that are consistent with observations; spatial patterns for Tkv, Cr, Brk and Pent that are consistent with observations; and levels of total Dpp inside cells that are much higher than outside (Kicheva et al., 2007; Zhou et al., 2012).

After identifying parameter sets that fulfilled these criteria we turned to the other mutant genotypes. For ubi-*tkv* (Tkv uniformly expressed in the entire disc), we take *tkv2*=*tkv3*=0, and adjust *tkv1* to obtain a final pMad gradient similar to that of wildtype (since that is what is observed experimentally). Similarly, for uniform *dally*, we take *cr2*=*cr3*=0, and adjust *cr1*. Parameters used for double mutant and triple mutants, combine several of these alterations. 10,000 random values of tkv1 for ubi-*tkv* and cr1 for uniform dally were explored and tested for quality of fit to the experimental data. Lastly, we locally perturbed all parameters over a four-fold range, exploring 500,000 parameters sets, to find parameter set used to produce the results in Figures 5J and 6. These parameters are listed in the table below, followed by the full simulation results for all genotypes.

Also given in the table below are the parameters *a*, *b* and *n* in the growth rate equation that was used for simulation (see Section 2, above). As shown in Fig. S3, homozygous *pent* mutant discs grow more slowly than wild type discs, and therefore are fit with a different value of parameter *b*. Although we did not produce full growth curves for all genotypes, the distribution of posterior compartment sizes that we observed suggest that all of the other genotypes that scale poorly (ubi-*tkv*, *pent*$^{-/-}$; uniform *dally, pent*$^{-/-}$*;* ubi-*tkv*, uniform *dally*; ubi-*tkv*, uniform *dally*, *pent*$^{-/-}$)

grow at a rate similar to *pent*$^{-/-}$, whereas the others grow at a similar rate to wildtype (i.e. *pent*$^{+/-}$; ubi-*tkv*; uniform *dally*). The value of *b* used in simulation was therefore selected accordingly from the wildtype and the *pent*$^{-/-}$ values.

**Wildtype Parameters:**

| Parameter | Value | Units | Reference |
|---|---|---|---|
| $D_{dpp}$ | 20 | μm$^2$*sec$^{-1}$ | (Zhou et al., 2012) |
| $k_1$ | $1.00\times10^5$ | M$^{-1}$*sec$^{-1}$ | (Lander et al., 2002) |
| $k_2$ | $3.68\times10^6$ | M$^{-1}$*sec$^{-1}$ | |
| $k_3$ | $2.34\times10^3$ | M$^{-1}$*sec$^{-1}$ | |
| $k_4$ | 1.26 | M$^{-1}$*sec$^{-1}$ | |
| $k_5$ | $4.47\times10^4$ | M$^{-1}$*sec$^{-1}$ | |
| $k_{r1}$ | $3.80\times10^{-3}$ | sec$^{-1}$ | |
| $k_{r2}$ | $8.30\times10^{-5}$ | sec$^{-1}$ | |
| $tkv1$ | $3.19\times10^{-12}$ | M *sec$^{-1}$ | |
| $tkv2$ | $2.87\times10^{-11}$ | M* sec$^{-1}$ | |
| $tkv3$ | $8.51\times10^{-12}$ | M *sec$^{-1}$ | |
| $cr1$ | $9.12\times10^{-12}$ | M *sec$^{-1}$ | |
| $cr2$ | $5.09\times10^{-11}$ | M *sec$^{-1}$ | |
| $cr3$ | $4.75\times10^{-11}$ | M* sec$^{-1}$ | |
| $v_{dpp}$ | $1.14\times10^{-11}$ | M *sec$^{-1}$ | |
| $v_{pmad}$ | $5.36\times10^{-2}$ | $EC_{brk}$*M$^{-1}$*sec$^{-1}$ | |
| $v_{pent}$ | $1.40\times10^{-3}$ | M* sec$^{-1}$ | |
| $v_{tf}$ | $1.53\times10^{-4}$ | $EC_{tkv}$*sec$^{-1}$ | |
| $v_{brk}$ | $5.39\times10^{-5}$ | $EC_{tf}$*sec$^{-1}$ | |
| $d_{tkv}$ | $9.10\times10^{-5}$ | sec$^{-1}$ | |
| $d_{cr}$ | $8.20\times10^{-5}$ | sec$^{-1}$ | |
| $d_{pmad}$ | $7.70\times10^{-4}$ | sec$^{-1}$ | |
| $d_{pent}$ | $7.10\times10^{-5}$ | sec$^{-1}$ | |
| $d_{tf}$ | $6.10\times10^{-5}$ | sec$^{-1}$ | |
| $d_{brk}$ | $4.10\times10^{-5}$ | sec$^{-1}$ | |
| $d_{dpp}$ | $1.00\times10^{-4}$ | sec$^{-1}$ | |

| | | | |
|---|---|---|---|
| $d_{dpptkv}$ | $4.00\times10^{-5}$ | sec$^{-1}$ | |
| $d_{dpptkv*}$ | $6.50\times10^{-5}$ | sec$^{-1}$ | |
| $d_{dppcr}$ | $7.80\times10^{-5}$ | sec$^{-1}$ | |
| $EC_{brk}$ | $2.80\times10^{-2}$ | -- | |
| $EC_{tkv}$ | 0.40 | -- | |
| $EC_{cr}$ | 1 | $EC_{tkv}$ | |
| $EC_{tf}$ | $7.50\times10^{-2}$ | -- | |
| $EC_{pent}$ | $1.14\times10^{1}$ | $EC_{brk}$ | |
| $p$ | 0.12 | -- | |
| $x_0$ | 0.1 | $\mu m$ | |
| $a$ | 0.1168 | sec$^{-1}$ | Figure S3 |
| $b$ | $4.77\times10^{-6}$ | -- | Figure S3 |
| $n$ | 3 | -- | Figure S3 |

**Parameter alterations in mutant genotypes:**

| Parameter | Value | Units | Reference |
|---|---|---|---|
| | ***pent*$^{+/-}$** | | |
| $v_{pent}$ | $7.00\times10^{-4}$ | M* sec$^{-1}$ | |
| | ***pent*$^{-/-}$** | | |
| $k_3$ | 0 | M$^{-1}$* sec$^{-1}$ | |
| $v_{pent}$ | 0 | M* sec$^{-1}$ | |
| $b$ | $6.02\times10^{-6}$ | -- | Figure S3 |
| | **ubi-*tkv*** | | |
| $tkv1$ | $4.26\times10^{-12}$ | M *sec$^{-1}$ | |
| $tkv2$ | 0 | M *sec$^{-1}$ | |
| $tkv3$ | 0 | M *sec$^{-1}$ | |
| | **uniform *dally*** | | |
| $cr1$ | $5.47\times10^{-11}$ | M *sec$^{-1}$ | |
| $cr2$ | $5.09\times10^{-11}$ | M *sec$^{-1}$ | |
| $cr3$ | $4.75\times10^{-11}$ | M* sec$^{-1}$ | |
| | **ubi-*tkv*, uniform *dally*** | | |
| $tkv1$ | $4.26\times10^{-12}$ | M *sec$^{-1}$ | |
| $tkv2$ | 0 | M *sec$^{-1}$ | |
| $tkv3$ | 0 | M *sec$^{-1}$ | |
| $cr1$ | $5.47\times10^{-11}$ | M *sec$^{-1}$ | |
| $cr2$ | 0 | M *sec$^{-1}$ | |

| | | | |
|---|---|---|---|
| *cr*3 | 0 | M* sec$^{-1}$ | |
| *b* | $6.02\times10^{-6}$ | -- | Figure S3 |
| **ubi-*tkv*, *pent*$^{-/-}$** | | | |
| *tkv*1 | $4.26\times10^{-12}$ | M *sec$^{-1}$ | |
| *tkv*2 | 0 | M *sec$^{-1}$ | |
| *tkv*3 | 0 | M *sec$^{-1}$ | |
| $k_3$ | 0 | M$^{-1}$* sec$^{-1}$ | |
| $v_{pent}$ | 0 | M* sec$^{-1}$ | |
| *b* | $6.02\times10^{-6}$ | -- | Figure S3 |
| **ubi-*tkv*, uniform *dally*, *pent*$^{-/-}$** | | | |
| *tkv*1 | $4.26\times10^{-12}$ | M *sec$^{-1}$ | |
| *tkv*2 | 0 | M *sec$^{-1}$ | |
| *tkv*3 | 0 | M *sec$^{-1}$ | |
| *cr*1 | $5.47\times10^{-11}$ | M *sec$^{-1}$ | |
| *cr*2 | 0 | M *sec$^{-1}$ | |
| *cr*3 | 0 | M* sec$^{-1}$ | |
| $k_3$ | 0 | M$^{-1}$* sec$^{-1}$ | |
| $v_{pent}$ | 0 | M* sec$^{-1}$ | |
| *b* | $6.02\times10^{-6}$ | -- | Figure S3 |

**wildtype**

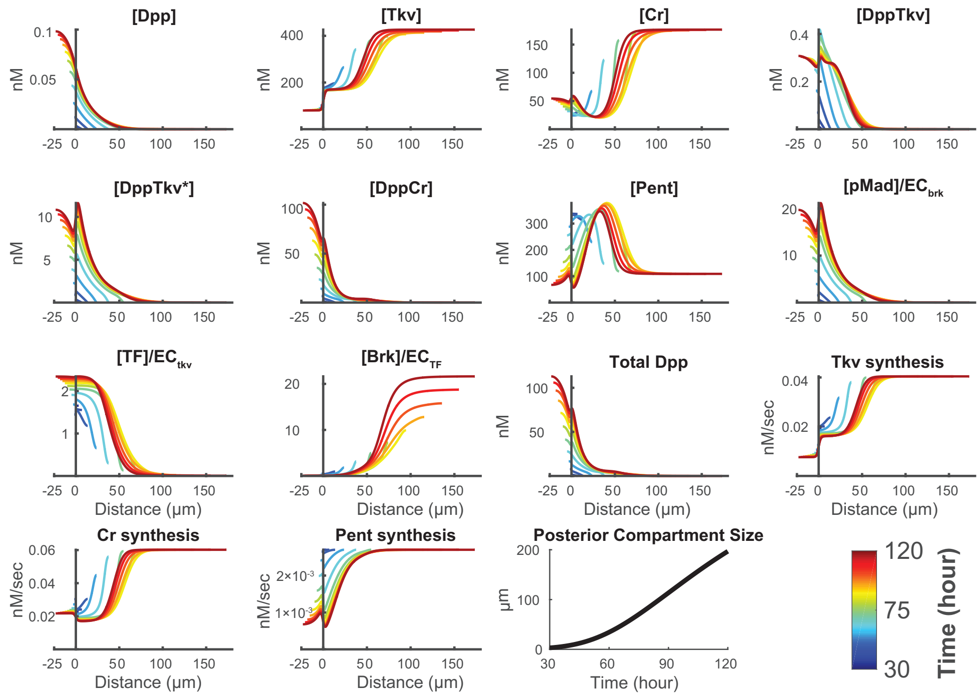

*pent*$^{+/-}$

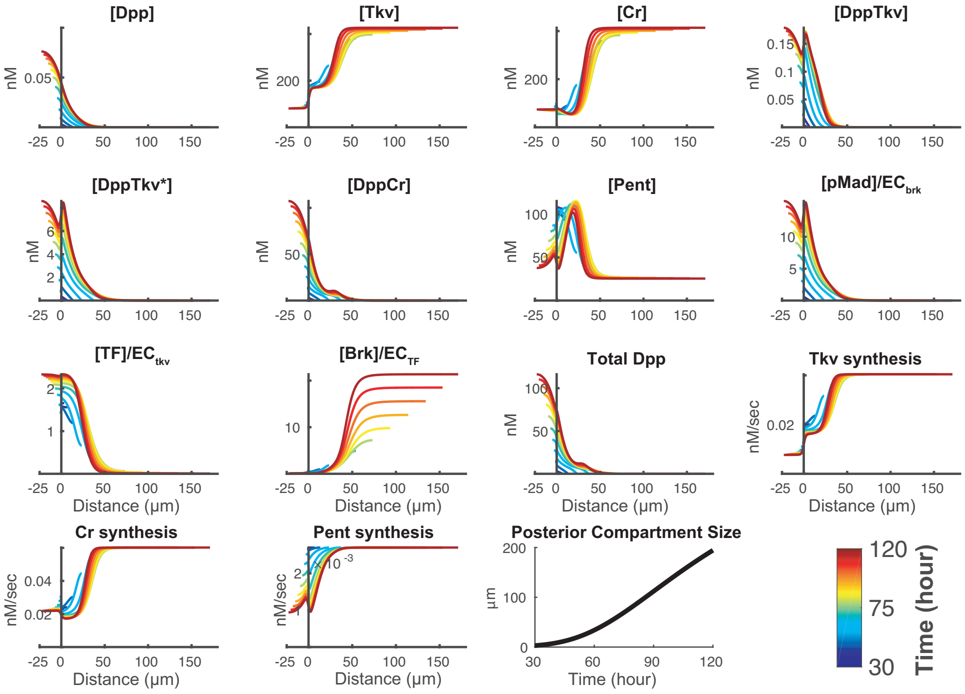
[Dpp]
[Tkv]
[Cr]
[DppTkv]
[DppTkv*]
[DppCr]
[Pent]
[pMad]/EC_brk
[TF]/EC_tkv
[Brk]/EC_TF
Total Dpp
Tkv synthesis
Cr synthesis
Pent synthesis
Posterior Compartment Size
nM
nM/sec
µm
Distance (µm)
Time (hour)
Time (hour)

*pent*$^{-/-}$

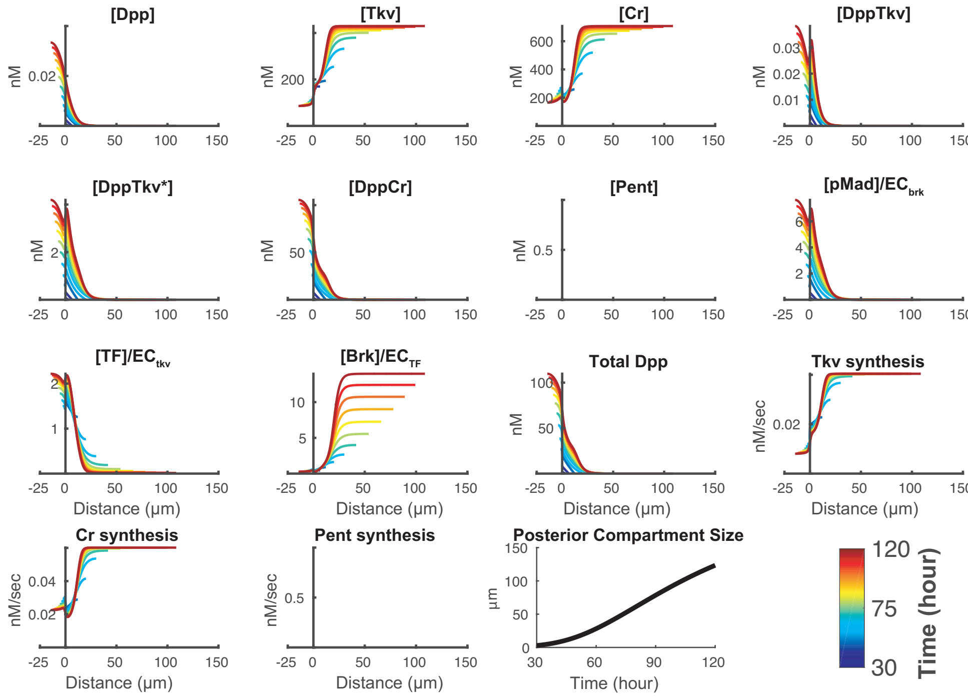
[Dpp]
[Tkv]
[Cr]
[DppTkv]
[DppTkv*]
[DppCr]
[Pent]
[pMad]/EC_brk
[TF]/EC_tkv
[Brk]/EC_TF
Total Dpp
Tkv synthesis
Cr synthesis
Pent synthesis
Posterior Compartment Size
nM
nM/sec
µm
Distance (µm)
Time (hour)
Time (hour)

### ubi-*tkv*

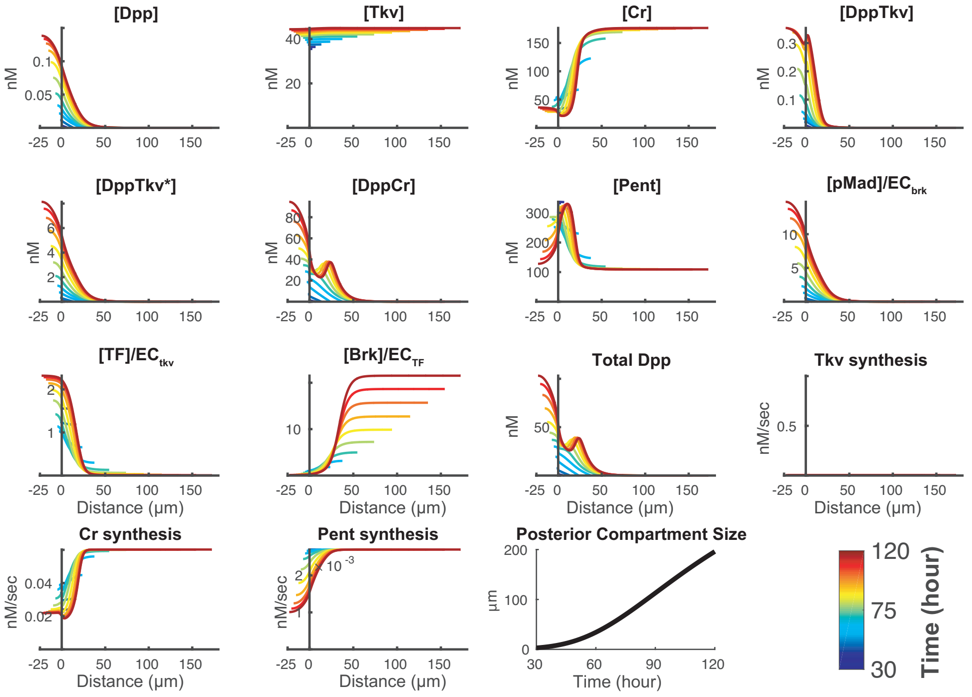


### uniform *dally*

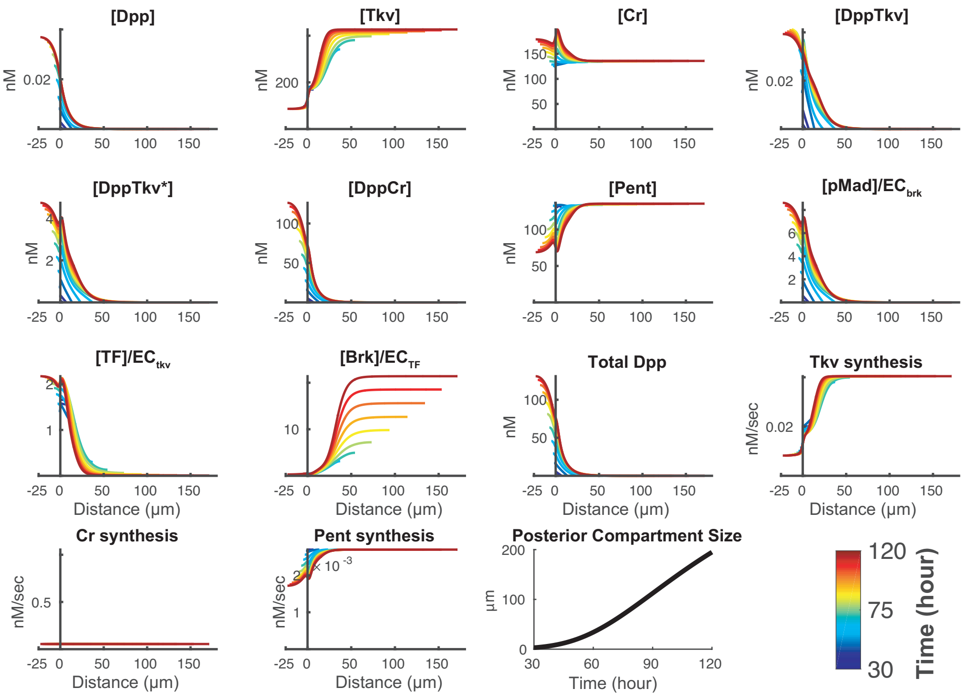

**ubi-*tkv*; *pent$^{-/-}$***

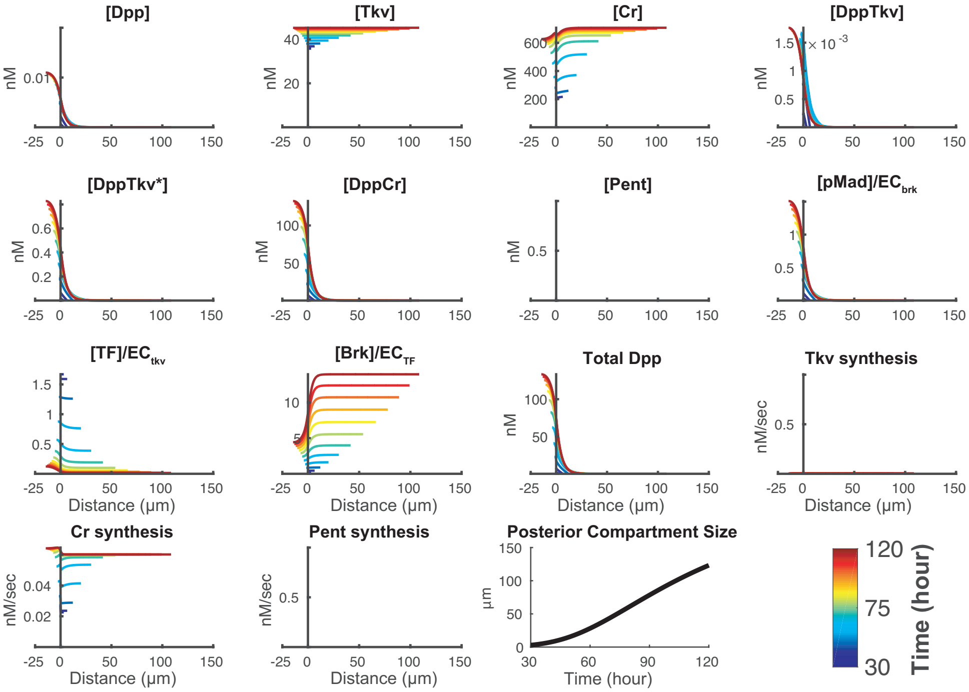


**ubi-*tkv*; uniform *dally***

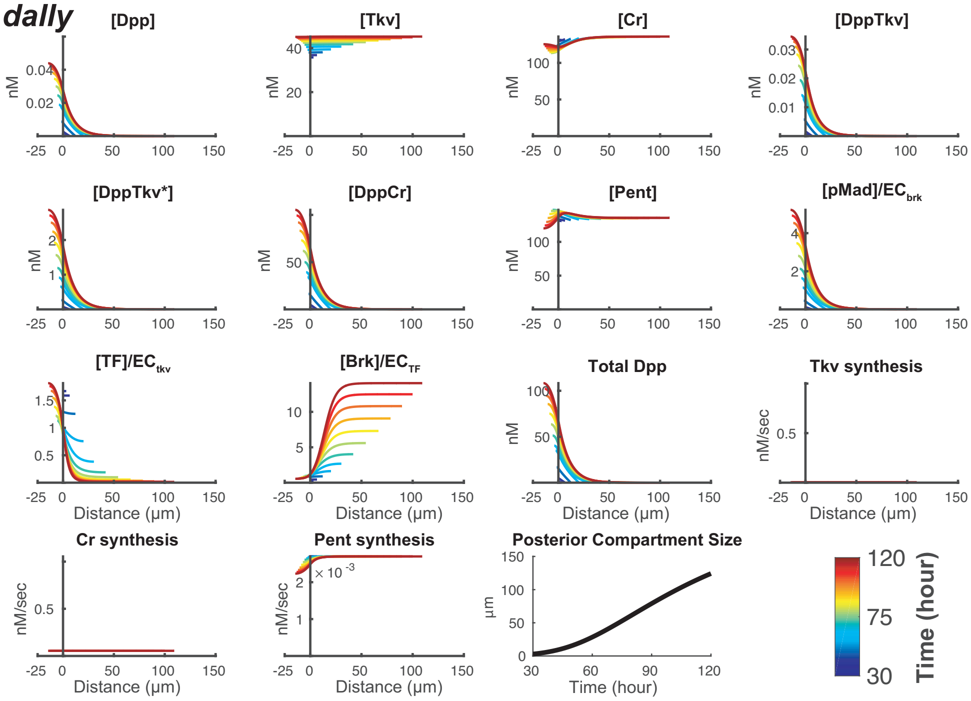

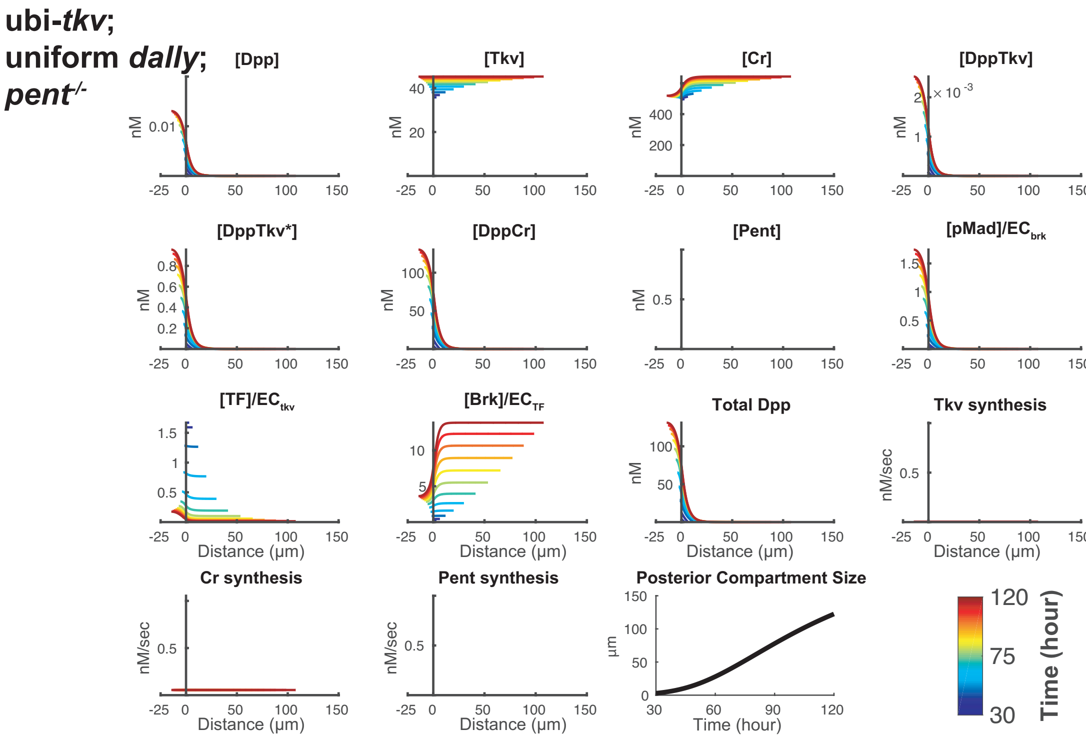


### Section 5. Measurement of Apparent decay length for pMad and intrinsic decay length of Dpp from simulations

To quantify the pMad apparent decay length ($\lambda_{app}$) in simulation data, we exclude the Dpp production region, and identify the location where pMad declines to 1/*e* of its maximum value. $\lambda_{app}$ is then defined as the distance between this location and the location where pMad is maximal.

We also quantify the intrinsic decay length ($\lambda_{\text{intrinsic}}$) of Dpp. The definition of intrinsic decay length of Dpp is given by

$$\lambda_{\text{intrinsic}} = \sqrt{\frac{D_{dpp}}{k_{\text{deg}}(x,t)}} \qquad (13)$$

where $D_{dpp}$ is the diffusion coefficient and $k_{\deg}(x,t)$ is the effective degradation rate constant at location $x$ and time $t$. The effective degradation rate constant is simply the sum of all of the terms on the right-hand side of the Dpp equation in (1), neglecting the diffusion term, divided by [Dpp]. Thus,

$$k_{\deg}(x,t) = k_1[Tkv] + k_2[Cr] + \frac{d_{dpp}}{1+\left(\frac{x}{prod(t)}\right)^{20}} - \frac{k_{r1}[DppTkv] - k_{r2}[DppCr]}{[Dpp]}. \qquad (14)$$

**Section 6. Local Sensitivity Analysis**

To determine whether the qualitative behaviors of the system are strongly dependent on the choice of parameters values, we systematically varied all parameters up and down by 10-fold, and measured the effect on $\lambda_{app}$ of pMad. Specifically, we calculated the ratio between $\lambda_{app}$ at the end of the simulation for the unperturbed case and the perturbed case. This was done for four genotypes (wildtype; $pent^{-/-}$; ubi-*tkv*; uniform *dally*) and the results are shown below. Most perturbations produced relatively small changes. The system is relatively insensitive to $k_4$, $k_5$, $k_{r1}$, $d_{dpp}$ and $d_{dpptkv}$, whereas it is relatively sensitive to $cr1$ and $v_{dpp}$.

| **Ratio of perturbed $\lambda_{app}$ to unperturbed $\lambda_{app}$.** | | | | | | | | |
|---|---|---|---|---|---|---|---|---|
| | wildtype | | $pent^{-/-}$ | | ubi-*tkv* | | uniform *dally* | |
| Fold perturbation: | x 0.1 | x 10 | x 0.1 | x 10 | x 0.1 | x 10 | x 0.1 | x 10 |
| $k_1$ | 0.9 | 0.5 | 0.5 | 0.8 | 0.6 | 0.8 | 0.8 | 0.5 |
| $k_2$ | 1.5 | 0.5 | 2.1 | 0.2 | 2.5 | 0.2 | 1.3 | 0.3 |
| $k_3$ | 0.7 | 1 | 1 | 1 | 0.4 | 1.4 | 0.7 | 1.2 |
| $k_4$ | 0.9 | 1 | 1.1 | 1 | 0.9 | 1 | 1 | 1 |
| $k_5$ | 1 | 0.9 | 1 | 1 | 1 | 1.1 | 1 | 0.9 |
| $k_{r1}$ | 1 | 0.9 | 1 | 1 | 1 | 1 | 1 | 1 |
| $k_{r2}$ | 0.8 | 1.4 | 0.6 | 1.8 | 0.4 | 2.2 | 0.7 | 1.2 |
| $tkv1$ | 1.1 | 0.7 | 1 | 0.8 | 0.6 | 0.8 | 1.1 | 0.5 |
| $tkv2$ | 1.1 | 0.6 | 0.6 | 0.8 | 1 | 1 | 0.8 | 0.6 |

| | | | | | | | | |
|---|---|---|---|---|---|---|---|---|
| $tkv3$ | 1.2 | 0.6 | 1 | 0.9 | 1 | 1 | 1.1 | 0.5 |
| $cr1$ | 1.2 | 0.3 | 1.2 | 0.4 | 1.5 | 0.3 | 1.7 | 0.2 |
| $cr2$ | 1.2 | 0.1 | 1.2 | 0.2 | 1.8 | 0.2 | 1 | 1 |
| $cr3$ | 1 | 1.2 | 1 | 0.7 | 1.1 | 0.5 | 1 | 1 |
| $v_{dpp}$ | 0.5 | 1.9 | 0.5 | 2.4 | 0.5 | 3.4 | 0.6 | 0.8 |
| $v_{pmad}$ | 0.5 | 0.6 | 0.4 | 0.9 | 0.6 | 0.9 | 0.7 | 0.5 |
| $v_{pent}$ | 0.6 | 1.2 | 1 | 1 | 0.3 | 1.7 | 0.5 | 1.4 |
| $v_{brk}$ | 1 | 0.8 | 0.9 | 0.5 | 1.5 | 0.6 | 0.7 | 0.7 |
| $d_{tkv}$ | 0.6 | 0.7 | 0.8 | 0.5 | 0.9 | 0.6 | 0.5 | 0.8 |
| $d_{cr}$ | 1 | 1.3 | 0.3 | 2.1 | 0.7 | 1.7 | 0.6 | 1.1 |
| $d_{pmad}$ | 0.7 | 0.5 | 1 | 0.4 | 1 | 0.6 | 0.5 | 0.7 |
| $d_{pent}$ | 1.1 | 0.7 | 1 | 1 | 1.7 | 0.4 | 1.1 | 0.7 |
| $d_{tf}$ | 0.7 | 1 | 0.7 | 1 | 0.6 | 1.1 | 0.7 | 1 |
| $d_{brk}$ | 1 | 1 | 0.7 | 0.9 | 0.6 | 1.5 | 0.9 | 0.7 |
| $d_{dpp}$ | 1 | 1 | 1 | 1 | 1 | 1 | 1 | 1 |
| $d_{dpptkv}$ | 1 | 0.9 | 1 | 1 | 1 | 1 | 1 | 1 |
| $d_{dpptkv*}$ | 0.9 | 0.5 | 1.1 | 0.4 | 1.4 | 0.5 | 0.7 | 0.7 |
| $d_{dppcr}$ | 1.4 | 0.9 | 1.6 | 0.5 | 2 | 0.4 | 1.2 | 0.7 |
| $EC_{brk}$ | 1 | 0.4 | 0.9 | 0.4 | 1.5 | 0.6 | 0.7 | 0.5 |
| $EC_{tkv}$ | 1.1 | 0.7 | 0.9 | 0.8 | 1 | 1 | 0.8 | 0.5 |
| $EC_{cr}$ | 1.2 | 0.8 | 1.3 | 0.7 | 1.8 | 0.6 | 1 | 1 |
| $EC_{tf}$ | 0.8 | 1 | 0.5 | 0.9 | 0.6 | 1.5 | 0.7 | 0.7 |
| $EC_{pent}$ | 0.6 | 1.4 | 1 | 1 | 0.5 | 1.3 | 0.5 | 0.9 |

#### Section 7. A reduced model that exhibits pseudo-source-sink behavior

We may illustrate the principle of pseudo-source-sink scaling using a much simpler model than (1). In this model, shown below, only three species are considered: ligand ([L]), receptor ([R]) and the complex between ligand and receptor ([LR]).

$$
\begin{cases}
\dfrac{d[L]}{dt} = D\Delta[L] + \begin{cases} v_L - d_L[L], \text{ if } x < 0.12x_{\max} \\ -k_{on}[L][R], \text{ if x} \geq 0.12\text{x}_{\max} \end{cases} \\
\dfrac{d[R]}{dt} = \dfrac{v_R}{1+\left(g[LR]\right)^2} - k_{on}[L][R] - d_R[R], \\
\dfrac{d[LR]}{dt} = k_{on}[L][R] - d_{LR}[LR].
\end{cases}
\quad (15)
$$

The spatial domain is $[0, x_{\max}]$. The effects of dilution and advection are neglected, as their impact on the full model turned out to be minimal (at least for the parameters chosen in Fig. 5J and 6). This enabled us to solve the system at steady state on a variety of fixed domain sizes, rather than model continuous domain growth. As in the full model, ligand is produced in a localized production region that grows proportionately with the rest of the disc. The morphogen diffuses and binds receptors, but here, dissociation from receptors is neglected as it is thought to be slow; the binding event may be understood as representing the combination of binding, uptake and destruction in a single step. Inside the morphogen production region, where we know that receptor and co-receptor levels are handled differently than elsewhere, we replace the usual receptor interaction term with a first order morphogen decay term $d_L[L]$, meant to represent the aggregate of those interactions within the production region. The production of receptor is subject to negative feedback from the amount of complex [*LR*] (which is taken to be a proxy for "signal" from the morphogen). Parameter *g* is the reciprocal of an $EC_{50}$, and it reflects the strength of feedback. Setting *g*=0 is equivalent to removing feedback.

We can simply non-dimensionalize this system to make both time and space unitless. The three species in (15) are thus re-named according to:

$$
\begin{cases}
\mu = \dfrac{k_{on}}{d_R}[L], \\
\rho = \dfrac{d_R}{v_R}[R], \\
\omega = \dfrac{d_R}{v_R}[LR].
\end{cases}
\quad (16)
$$

For receptor [R] and the complex [LR], this transformation is equivalent to normalization to the level of free receptor that would obtain in the absence of any feedback or ligand, $R_{\max} = \frac{v_R}{d_R}$. We also nondimensionalize space by defining the unit of distance to be $\lambda_0$, the intrinsic decay length that would be observed in the absence of ligand binding or feedback:

$$\lambda_0 = \sqrt{\frac{D}{k_{on} R_{\max}}} = \sqrt{\frac{D d_R}{k_{on} v_R}}. \tag{16}$$

Finally, we may nondimensionalize time by scaling it to the inverse of the degradation rate of receptor $d_R$ (although the time scale is not relevant to the steady state analysis of (15), it simplifies numerical solution by time-evolution). Thus, the transformation from original coordinates $(x,t)$ to the new coordinates $(X,\tau)$ is given by:

$$\begin{cases} X = \dfrac{x}{\lambda_0}, \\ \tau = t d_R. \end{cases} \tag{17}$$

The nondimensionalized equations are therefore given by

$$\begin{cases} \dfrac{d\mu(X,\tau)}{d\tau} = k\Delta\mu(X,\tau) + \begin{cases} -k\varphi\mu(X,\tau) + k\upsilon, & \text{if } X < 0.12X_{\max} \\ -k\mu(X,\tau)\rho(X,\tau), & \text{if } X \geq 0.12X_{\max} \end{cases}, \\ \dfrac{d\rho(X,\tau)}{d\tau} = \dfrac{1}{1+\left(\gamma\omega(X,\tau)\right)^2} - \mu(X,\tau)\rho(X,\tau) - \rho(X,\tau), \\ \dfrac{d\omega(X,\tau)}{d\tau} = \mu(X,\tau)\rho(X,\tau) - \xi\omega(X,\tau). \end{cases} \tag{18}$$

The five nondimensional free parameters in (18) are related to the parameters in (15) according to:

$$
\begin{cases}
k = \dfrac{k_{on} v_R}{d_R^2}, \\
\varphi = \dfrac{d_L d_R}{k_{on} v_R} \\
\upsilon = \dfrac{v_L}{v_R}, \\
\xi = \dfrac{d_{LR}}{d_R}, \\
\gamma = g \dfrac{v_R}{d_R}.
\end{cases}
\tag{19}
$$

In addition, $X_{\max}$, the spatial size of the domain scaled to $\lambda_0$, enters as a sixth parameter that is required to specify the boundary condition opposite the production region. At the start of the production region we impose a no-flux boundary condition, to reflect the spatial symmetry of the system. At the end of the gradient region, $x$=$x_{\max}$, we impose an absorbing boundary condition.

Below we show the steady-state shapes of a series of eight gradients associated on increasing compartment sizes. Curves are color-coded to represent increasing domain sizes (which, in this case were $X_{\max}$ = 1, 2, 3, 4, 5, 6, 9, and 12. To obtain the results labeled "No Feedback", we set the feedback strength $\gamma$ to zero, and adjusted the morphogen production rate to match so that the results for LR near the origin would be similar in the two cases.

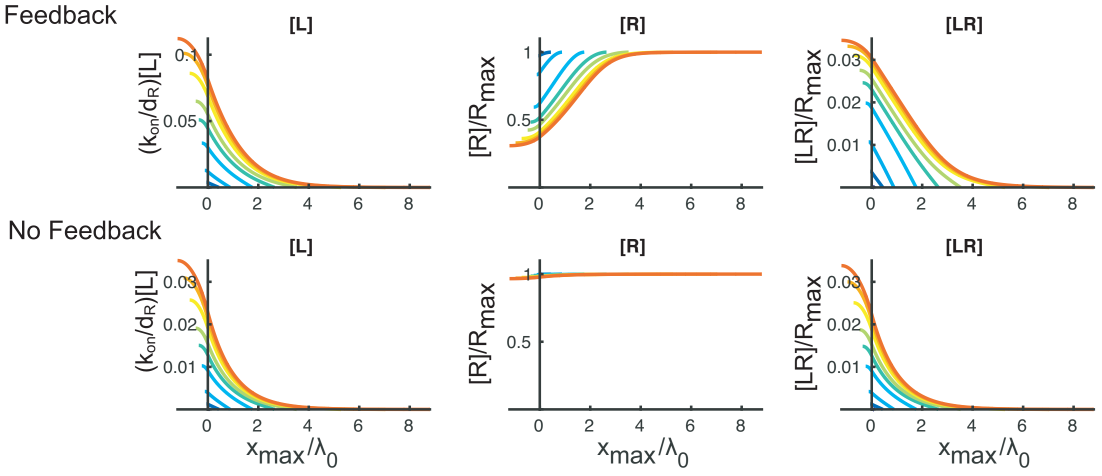

The third panel of each of these cases is reproduced in the main Figures 6C-D, and shows how feedback enables gradients to remain quasi-linear, and continue scaling, for much longer. Note also the growing suppression of receptor expression in the Feedback case. The parameters used were

| Parameter | with feedback | no feedback |
|---|---|---|
| $k$ | 1 | 1 |
| $\varphi$ | 1 | 1 |
| $\upsilon$ | 0.15 | 0.05 |
| $\xi$ | 1 | 1 |
| $\gamma$ | 40 | 0 |

To more thoroughly understand the behavior of the reduced model, we explored a large number of random parameters. It is clear, from (18), that the parameter $k$ drops out in the steady state, so that in any exploration of parameters in which we are only interested in steady-state behavior we can simply fix $k$ to be 1. We then randomly generated $\varphi$, $v$, $\xi$ and $X_{max}$ using Latin hypercube sampling, initially running simulations with no feedback ($\gamma$=0). A total simulation time of T=10,000 allowed us to obtain a steady-state solution numerically. The numerical steady-state solution of ligand-receptor complex is denoted by $\omega(X)_{SS}^{noFB}$. Next, we ran simulations with feedback. Rather than choose values of $\gamma$ completely at random, we selected them so as to exclude those that would provide only trivial amounts of feedback, as well as those that would provide so much feedback that receptors would be fully suppressed from the start. In particular, we chose $\gamma$ as defined by

$$\gamma = \frac{1}{0.25 \max\limits_{X \in [0, X_{max}]} \omega(X)_{SS}^{noFB}}. \qquad (20)$$

The parameters ranges that were explored are shown below

| Parameter | Range |
|---|---|
| $\varphi$ | $(10^{-2}, 10^{2})$ |
| $\upsilon$ | $(10^{-2}, 10^{2})$ |
| $\xi$ | $(10^{-2}, 10^{2})$ |
| $X_{max}$ | (0.1,10) |

The results of comparing feedback and no-feedback scenarios for 1000 randomly generated parameter sets are shown below. They are plotted as in Fig 6E, with the apparent decay length $\lambda_{app}$ (normalized to $\lambda_0$, where $\lambda_0$ is effectively equivalent to $\lambda_{intrinsic}$ in the absence of feedback; note the logarithmic axis) being plotted against the field size (normalized to $\lambda_0$).

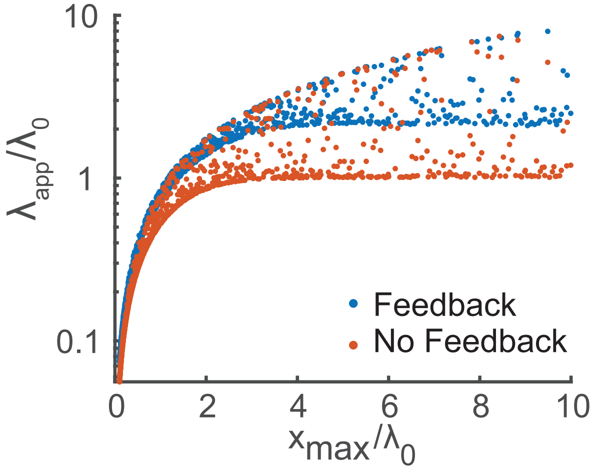


Note the ability of feedback to produce values of $\lambda_{app}$ much greater than observed without feedback. This effect is even more apparent if we eliminate those cases in which receptor saturation *S*—defined as the fraction of total receptors that are occupied (i.e. $S = [LR]/([LR]+[R])$)--exceeds 50% at the origin (the boundary between the production region and the rest of the domain), shown below.

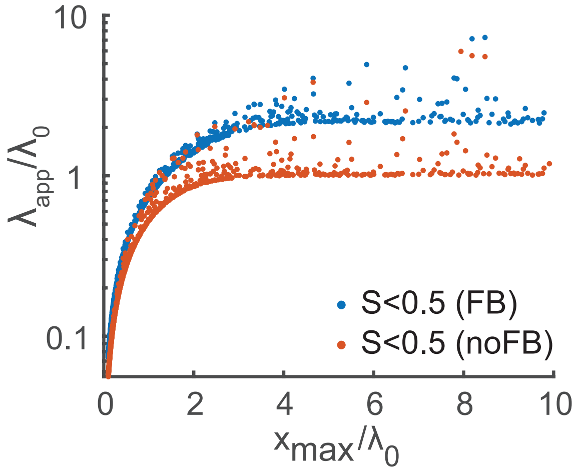


The reason for this is that morphogen decay is a function of *free* receptor level, and saturation amounts to lowering free receptor level. Thus, a gradient can, in principle, extend its apparent decay length simply by saturating receptors, but there are two reasons why this regime is likely to be un-biological. First, in this regime gradients adopt sigmoidal shapes, and such shapes are not observed in any known morphogen system. Second, a consequence of operating in this regime is that gradient position becomes extremely sensitive to small changes in the rate of morphogen production (Lander et al., 2009).

The above two plots do not do a good job of comparing individual parameter sets—with and without feedback—against each other. To do that, we replot the data as follows. The abscissa gives the apparent decay lengths, relative to the intrinsic decay length $\lambda_0$, for cases without feedback $\lambda_{app}^{noFB}$. The ordinate shows the ratio between $\lambda_{app}$ when feedback is present, and $\lambda_{app}$ without feedback. Each dot represents a single parameter set. Three colors are used: blue indicates that receptor saturation is below 50% at the origin for both the feedback scenario and the no-feedback scenario; red indicates that saturation is below 50% for the no-feedback case but not the no-feedback case; yellow indicates the saturation if above 50% in both cases (there were no parameter sets for which the feedback case was less than 50% saturated and the no-feedback case was not).

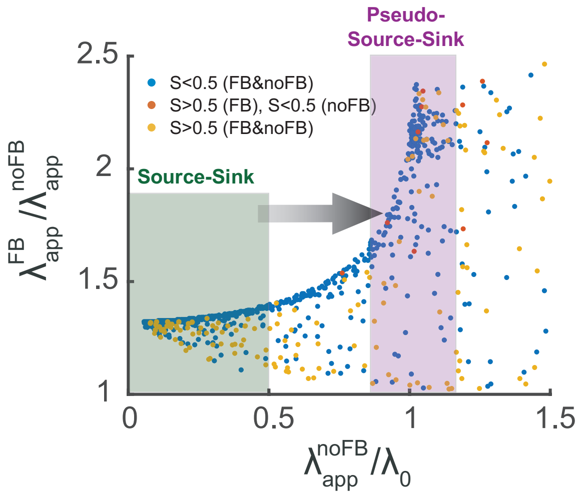


Notice that, when $\lambda_{app}^{noFB}$ is less than half the value of $\lambda_0$, the improvement in $\lambda_{app}$ that comes from feedback is always modest. This is because, in this regime, gradient shape is close to linear even in the absence of feedback, and thus both the feedback and non-feedback case display true source-sink scaling.

However, once $\lambda_{app}^{noFB}$ is on the order of $\lambda_0$ or larger, the improvement in $\lambda_{app}$ due to feedback is much greater for almost all parameter sets: this is the pseudo-source-sink scaling regime, in which gradients remain quasilinear and scale automatically, even though the true sink is located many values of $\lambda_{intrinsic}$ away. In highly saturated regimes (red and yellow symbols), however, some of the feedback cases perform no better than the no-feedback cases, presumably because scaling due to saturation of receptors does not require feedback to occur.
